# Sub-Millivolt Voltage Imaging Reveals Gap Junction-Mediated Bioelectric Contact Inhibition

**DOI:** 10.64898/2026.02.10.701308

**Authors:** Philipp Rühl, Rama Hussein, Stefanie Reuter, Konrad Frahnert, Anagha G. Nair, Ralf Mrowka, Roland Schönherr, Stefan H. Heinemann

**Affiliations:** Center for Molecular Biomedicine, Department of Biophysics, Friedrich Schiller University Jena and Jena University Hospital, D-07745, Jena, Germany; ThIMEDOP, Jena University Hospital, Am Klinikum 1, D-07747 Jena, Germany; Experimental Nephrology, Clinics for Internal Medicine III, Jena University Hospital, Am Klinikum 1, D-07747 Jena, Germany

## Abstract

Sub-millivolt membrane potential (*V*_m_) dynamics in multicellular non-excitable networks have remained largely invisible due to a lack of sufficiently sensitive imaging tools. Here, we introduce rEstus2s, a next-generation genetically encoded voltage indicator that overcomes this barrier by enabling high-resolution *V*_m_ imaging across the full physiological resting *V*_m_ range (-100 to 0 mV). Using rEstus2s, we uncover bioelectric contact inhibition (BCI), a fundamental biophysical principle where gap junction coupling acts as a passive noise filter to stabilize *V*_m_. We demonstrate that the time-dependent variance of *V*_m_ (electrical volatility) relative to the number of cells (*n*) in a network follows a 1/*n* scaling law, reflecting a transition from stochastic single-cell behavior to collective electrical stability. While Ca^2+^-activated oncogenic ion channels, including ANO1 and K_Ca_3.1, promote pronounced electrical volatility in isolated cells, BCI effectively attenuates volatility in electrically coupled networks. Disruption of gap junction coupling in cancer cells abolishes BCI and restores high electrical volatility. These findings establish a unifying biophysical framework for understanding how multicellularity maintains electrical homeostasis in health and disease.

**Graphical Abstract:** 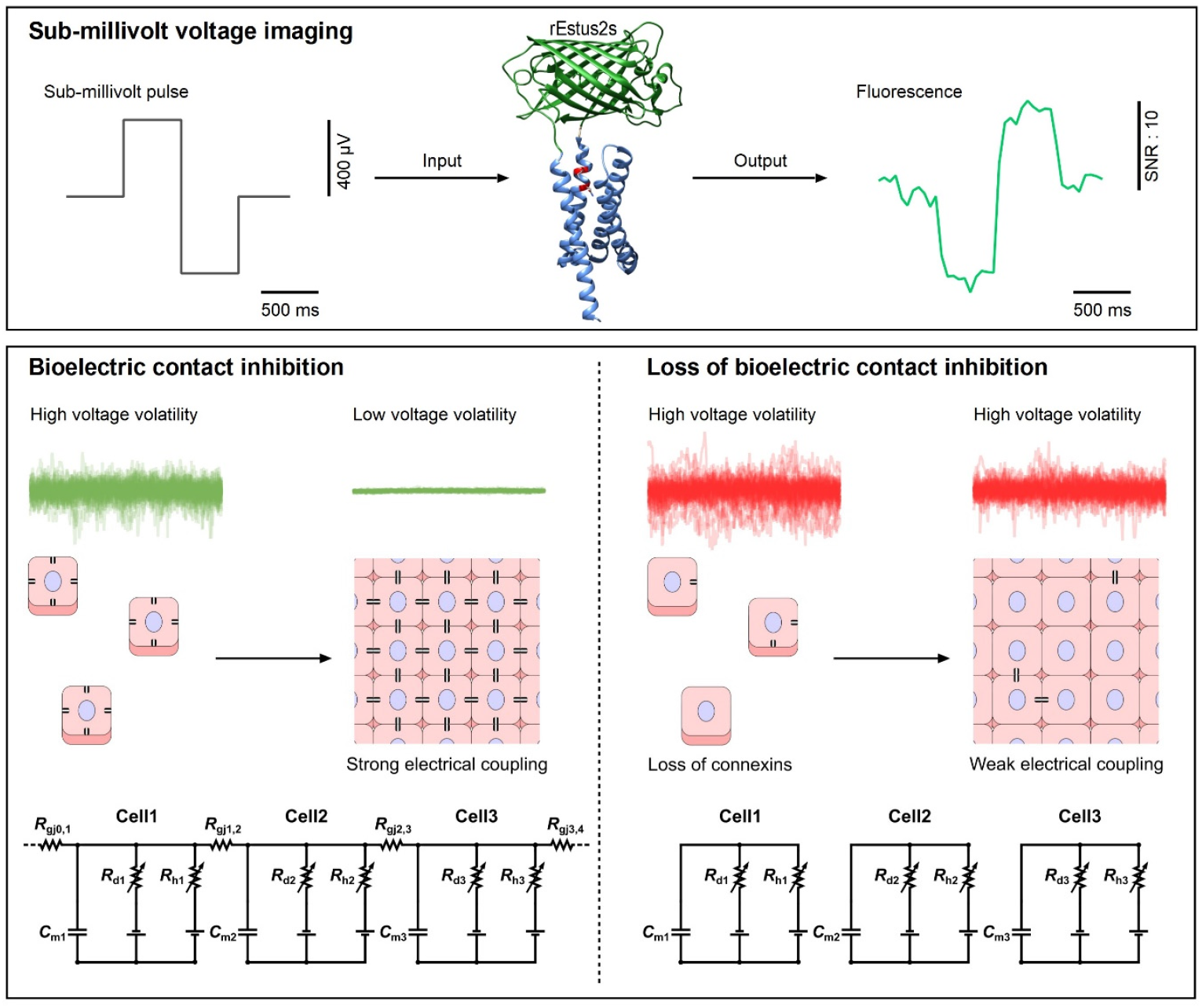

## Introduction

Bioelectric signaling outside the classical framework of excitable tissues is an emerging frontier in cell biology.^1–4^ Ion transport proteins play essential roles in development, tissue homeostasis, and tumorigenesis, yet the mechanistic pathways by which they influence these phenotypes remain poorly understood.^5^ In non-excitable cells, membrane potential (*V*_m_) has traditionally been viewed as a static property reflecting the balance of ion conductances. However, recent evidence supports an active regulatory role for *V*_m_ in non-excitable cells, as membrane depolarization has been shown to directly activate the ERK signaling pathway, a central driver of proliferation and oncogenic transformation.^6,7^ *V*_m_ also influences intracellular Ca^2+^ dynamics, either by regulating voltage-gated Ca^2+^ channels or by altering the electrochemical driving force for Ca^2+^ influx.^8,9^ Specific ion channels, often referred to as oncochannels, are implicated in nearly all hallmarks of cancer, including sustained proliferation, resistance to apoptosis, and invasion.^5,10–12^ Ca^2+^-activated ion channels such as ANO1 (TMEM16A, conducting Cl^−^) and K_Ca_3.1 (*KCNN4*, conducting K^+^) are frequently overexpressed in malignancies including head and neck squamous cell carcinoma (HNSCC) and pancreatic ductal adenocarcinoma (PDAC).^13–20^ Their upregulation stimulates ERK signaling,^14,21^ correlates with tumor size, and is associated with poor clinical prognosis.^16,21,22^

In contrast, connexins, which form gap junctions between neighboring cells, are widely regarded as tumor suppressors.^23–25^ Loss of gap junction coupling is a common early event in tumorigenesis, although connexins may reappear in advanced tumors where they promote migration and invasion.^24,26,27^ Mouse models of PDAC expressing assembly-deficient connexin 43 (Cx43) exhibit accelerated tumor onset and enhanced ERK activation.^27,28^ In non-excitable tissues, gap junctions are predominantly viewed as mediators of metabolite and second-messenger transfer or as structural scaffolds. Their electrical coupling function, central in excitable cells, is rarely addressed.^24^ Levin and colleagues ^1^ proposed that the loss of gap junctions in cancer marks a transition from altruistic, system-level behavior to an individualistic single-cell state, mediated largely through bioelectricity. In this framework, ion channels are thought to modulate long-term steady-state *V*_m_, shaping cellular phenotypes through sustained depolarization or hyperpolarization.^3,29,30^ However, recent studies, including our own, have revealed that many non-excitable cells also generate rapid, spontaneous *V*_m_ excursions on the second-to-millisecond timescale.^31–33^ Thus, it is plausible that ion channels and connexins play roles in shaping the *V*_m_ dynamics in multicellular networks of non-excitable cells. In a previous study, we observed that endogenous *V*_m_ fluctuations in HEK293T cells diminish as cells reach confluency.^31^ We propose that electrical coupling via connexins integrates individual cells into a syncytium that acts as a biophysical filter, attenuating *V*_m_ dynamics.^34,35^ Testing this hypothesis requires the ability to resolve subtle *V*_m_ fluctuations across the full physiological resting *V*_m_ range in intact, multicellular networks.

Conventional electrode-based electrophysiological techniques are sensitive but highly invasive and of low-throughput. Genetically encoded voltage indicators (GEVIs) offer a non-invasive alternative, converting electrical signals into optical readouts with high temporal resolution.^36–40^ GEVI development has largely focused on maximizing speed to capture neuronal action potentials, exemplified by indicators such as ASAP5 and JEDI-1P.^38,41^ For non-excitable cells, sensitivity rather than speed is the limiting factor.^31,32^ We previously addressed this challenge by developing rEstus, a bright and sensitive GEVI capable of resolving endogenous *V*_m_ dynamics in non-excitable cells.^31,42^ However, rEstus exhibits a steep response curve in the depolarized range (0 to -50 mV), resulting in signal saturation under hyperpolarized conditions.

Here, we present rEstus2s, a next-generation GEVI generated through rational engineering that provides a two-fold increase in *V*_m_ sensitivity over its precursor rEstus and enables detection of sub-millivolt *V*_m_ changes in the range of -100 to 0 mV. Using rEstus2s in combination with CRISPR-Cas9 connexin knockout models, we demonstrate that the ion channels ANO1 and K_Ca_3.1 enhance spontaneous *V*_m_ fluctuations in non-excitable cells, whereas cell-cell contact leads to a robust, cluster-size-dependent suppression of *V*_m_ fluctuations. This suppression, which we term bioelectric contact inhibition (BCI), strictly depends on electrical coupling through connexins. Together, our results establish BCI as a fundamental biophysical constraint on multicellular networks, providing new insight into *V*_m_ dynamics in health and disease.

## Results

### Development of rEstus2s for sub-mV imaging in the range of physiological resting *V*_m_

To obtain a GEVI with enhanced *V*_m_ sensitivity, we introduced mutations into the previously developed ultrasensitive GEVI, rEstus.^31^ rEstus comprises a voltage-sensing domain (VSD) fused to a circularly permuted EGFP (Fig. 1a). For clarity, all mutations in the cpEGFP part of rEstus are referenced according to the numbering in EGFP. The chromophore in rEstus is an EGFP-type TYG motif (Fig. 1b).^43^ Conformational changes in many cpEGFP-derived sensors are thought to affect the flexibility of the chromophore, thereby influencing the brightness of the sensor. Thus, we introduced the T65G mutation (T311G in rEstus), resulting in a GYG-type chromophore, which has been shown to enhance flexibility in the excited state in EGFP.^43^ The properties of rEstus and rEstus-T65G were characterized by the expression of N-terminal mKate2 fusion constructs of both sensors in HEK293T cells and subsequent fluorescence recordings under whole-cell voltage clamp conditions (Figs. 1c-f). mKate2 served as a reference to correct for fluorescence variations arising from differences in expression levels, as previously described.^31,44,45^ Strikingly, although the maximum brightness *(B*_max_) of rEstus-T65G was approximately one-third of that of rEstus, the overall *V*_m_ sensitivity of the fluorescence, characterized by the total fractional change in brightness (*B*_max_/*B*_min_), increased from 4.5-fold to 9.6-fold (Figs. 1d-f).

**Figure 1.**
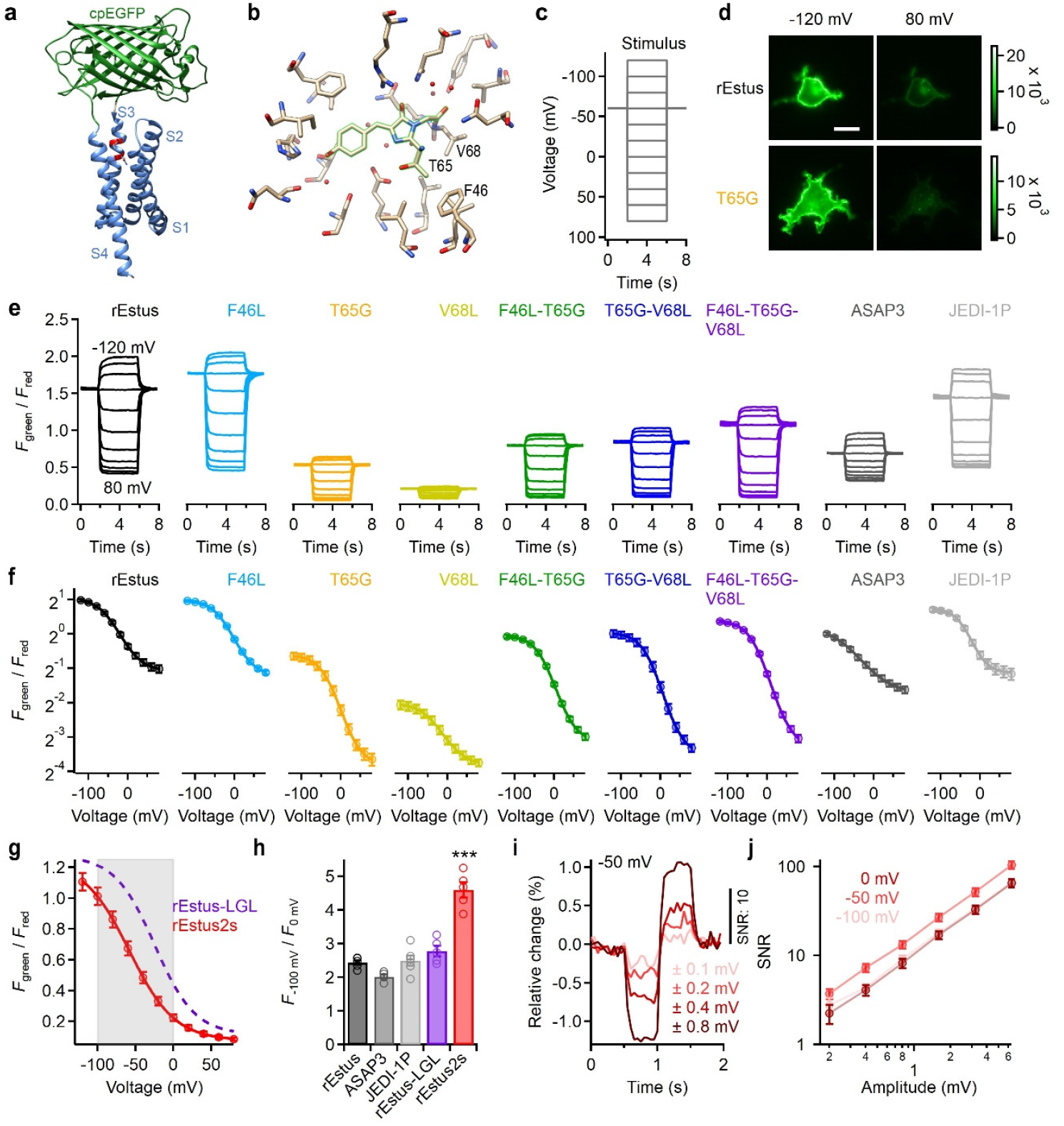
rEstus2s, a genetically encoded voltage indicator with enhanced voltage sensitivity. **a**, 3D model of rEstus2s; positions 138 and 141 are highlighted in red. **b**, Chromophore (green shading) and surrounding residues of EGFP (PDB: 2Y0G).^60^ **c**, Voltage pulse protocol. **d**, Fluorescence images (*F*_green_) of HEK293T cells expressing rEstus or rEstus-T311G (T65G in EGFP), voltage-clamped to -120 or 80 mV. Scale bar, 20 µm. **e**, Superposition of representative traces of *F*_green_ divided by the fluorescence of mKate2 (*F*_red_), which was fused to the N-terminus to account for variations in expression levels. Mutations are given with respect to EGFP. Results of the established GEVIs ASAP3 and JEDI-1P, are shown for comparison. **f**, *F*_green_ / *F*_red_ as a function of voltage (R-V) from the data shown in e. Data are means ± SEM, with superimposed Boltzmann-type fits (Equation 1). **g**, R-V relationship of rEstus2s; the fit to rEstus-LGL data is depicted for reference. The gray area indicates the physiological resting *V*_m_ range. **h**, Fold change in fluorescence across the physiological resting *V*_m_ range for the indicated GEVIs. **i**, Normalized fluorescence response of rEstus2s (without mKate2) to bipolar voltage steps with sub-mV amplitude from a holding *V*_m_ of -50 mV. Respective images were acquired at 20 Hz with 400/470 nm co-illumination. **j**, Signal-to-noise ratio (from i), calculated by dividing the steady-state fluorescence amplitude by the noise (standard deviation) at the holding *V*_m_ as a function of the stimulating voltage step size. Traces were recorded from holding voltages of 0 mV, -50 mV, and -100 mV. Data are means ± SEM, with n = 5.

To restore sensor brightness, we examined mutations that were introduced in other fluorescent proteins in combination with T65G. For example, EYFP (enhanced yellow fluorescent protein, with a GYG chromophore), harbors the additional mutations V68L, S72A, and T203Y, which are all near the chromophore (Fig. 1b).^46^ rEstus already contains S72A (S318A in rEstus), which we have previously shown to increase brightness relative to its precursor, ASAP3 by enhancing pH stability.^31^ Mutation V68L (V314L in rEstus) not only enhanced the brightness of rEstus-T65G but also retained its high *V*_m_ sensitivity (11.7 ± 0.5-fold, Figs. 1e,f). In contrast, the introduction of V68L into rEstus strongly reduced its maximal brightness and voltage sensitivity (Figs. 1e,f). T203Y facilitates π-stacking and shifts the excitation and emission spectrum in YFP-type proteins. In rEstus-T65G-V68L, the introduction of T203Y (T206Y in rEstus) severely reduced both brightness and voltage sensitivity (Supplementary Fig. S1). We further evaluated mutation F46L (F292L in rEstus), which was initially reported in the SEYFP-derived mVenus and enhances chromophore maturation at 37°C.^46^ In rEstus-T65G and rEstus-T65G-V68L, mutation F46L increased the brightness of the sensors, while the same mutation did not affect the brightness of rEstus. The triple mutant rEstus-F46L-T65G-V68L (rEstus-LGL) exhibited approximately twice the brightness of rEstus-T65G and approximately 65% of the brightness of rEstus.

While rEstus-LGL exhibited enhanced voltage sensitivity (Figs. 1e,f), the half-maximal activation voltage (*V*_half_) was shifted to -25 ± 3 mV, compared to -41 ± 3 mV in rEstus. Therefore, this mutant is best suited for cells with relatively depolarized *V*_m_. As we aimed for a sensor operating best at more negative *V*_m_, we introduced a previously characterized double mutation (G138N, T141I) into rEstus-LGL;^31,47^ both residues are located in the S3 helix of the VSD and face the outer vestibule (Fig. 1a). The resulting rEstus-LGL-G138N-T141I (rEstus-NILGL) had a left-shifted *V*_half_ of -58.3 ± 1.3 mV (Fig. 1g) and exhibited a relative brightness change between 0 to -100 mV of 4.60 ± 0.23-fold, which was greater than that of any of the other examined sensors (Fig. 1h). The voltage sensitivity (d*F*/*F* d*V*^-1^) of GEVIs is itself voltage dependent and can be described as the relative change in fluorescence (d*F*/*F*) in response to small voltage steps (d*V*). Because d*F*/*F* d*V*^-1^ was higher for rEstus-NILGL than for any of the other tested sensors within the physiological voltage range (Supplementary Fig. S2), we termed this new variant rEstus2s.

rEstus2s fluorescence responded to depolarizing steps in a biexponential time course. For a depolarizing step from -100 to 0 mV at 23°C, the time constants were 4.3 ± 0.4 ms (54.1 ± 1.6%) and 20.9 ± 2.0 ms (Supplementary Fig. S3). To estimate the effective detection limit of rEstus2s with our setup, we recorded square voltage pulses with amplitudes reaching into the sub-mV range at an acquisition rate of 20 Hz. While rEstus2s retains the standard green emission profile, it notably lacks the secondary excitation shoulder at approximately 400 nm observed in rEstus (Supplementary Fig. S4). Nevertheless, co-illumination of cells with 400 nm light allowed us to accelerate the kinetics of rEstus2s (Supplementary Fig. S5).^48^ Since voltage sensitivity is itself voltage-dependent, we recorded fluorescence-voltage relationships around three physiologically relevant resting voltages: 0, -50, and - 100 mV. At all holding voltages, voltage pulses in the sub-mV range were clearly detectable as fluorescence changes (Fig. 1i, Supplementary Fig. S6). At each holding voltage, a 500 ms square pulse with an amplitude of 400 µV gave signals with a signal-to-noise ratio greater than 3 (Fig. 1j). The highest SNR was reached at a holding *V*_m_ of -50 mV.

### Ca^2+^-activated K^+^ and Cl^-^ ion channels drive *V*_m_ volatility in non-excitable cells

Leveraging the enhanced voltage sensitivity of rEstus2s, we investigated how the Ca^2+^-activated oncochannels ANO1 (Cl^-^) and K_Ca_3.1 (K^+^) influence the *V*_m_ dynamics of non-excitable HEK293T cells (Fig. 2a). Given their respective ion selectivities, ANO1 and K_Ca_3.1 are expected to generate electrical currents determined by the Nernst potentials for Cl^-^ and K^+^ within the HEK293T system. By co-encoding the ion channel genes and rEstus2s on a single plasmid, we ensured consistent expression of both sensor and channels. Stable expression and function of ANO1 (Fig. 2b), K_Ca_3.1 (Fig. 2c), and rEstus2s (Fig. 2d) were subsequently validated using whole-cell patch-clamp and fluorescence imaging.

**Figure 2.**
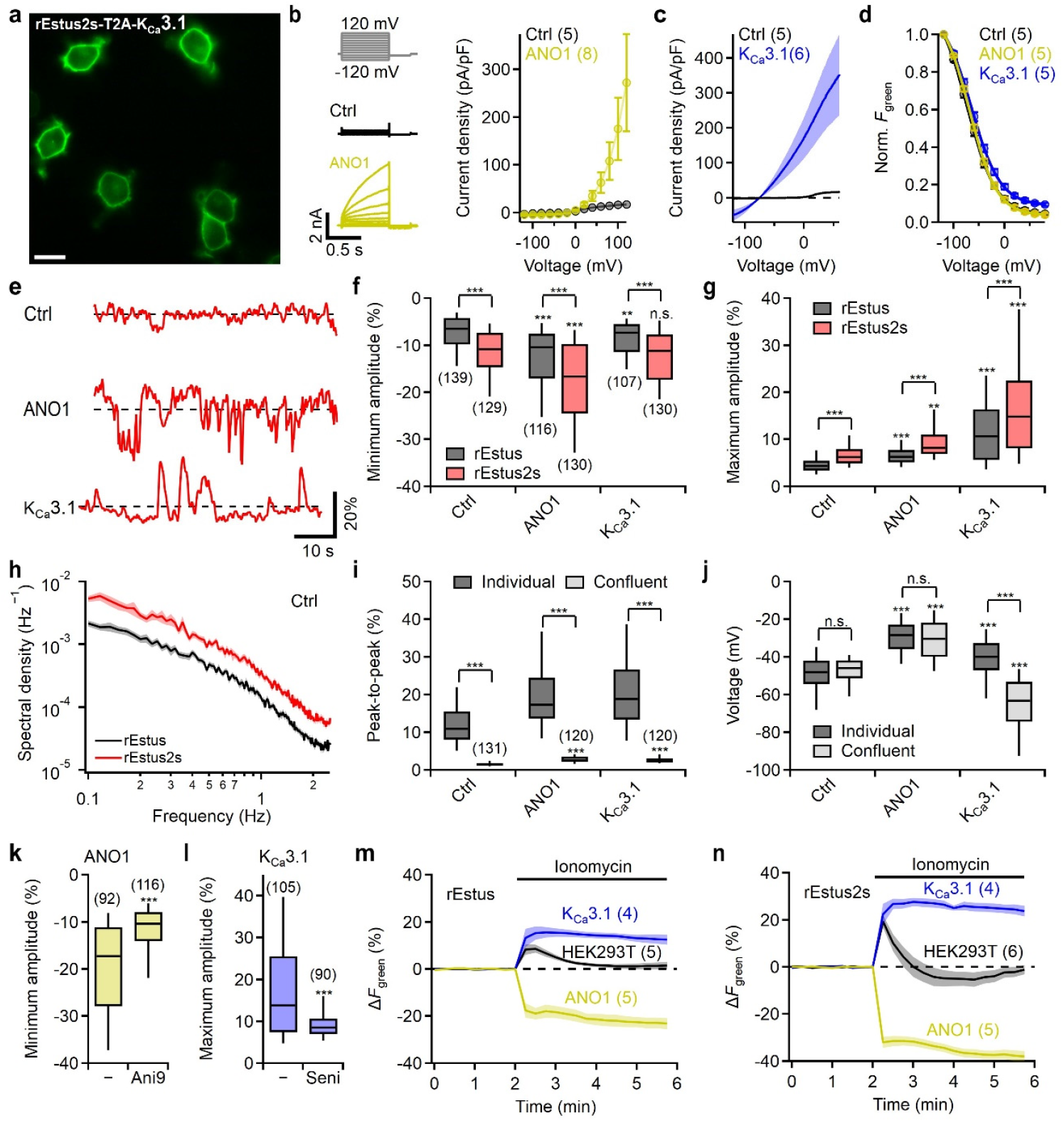
Ca^2+^-activated K^+^ and Cl^−^ channels enhance *V*_m_ volatility in HEK293T cells. **a**, Fluorescence image of HEK293T cells stably expressing rEstus2s-T2A-K_Ca_3.1. Scale bar, 20 µm. **b**, Whole-cell patch-clamp validation of ANO1 expression. Left: Representative current traces elicited by voltage steps from -120 mV through 120 mV. Right: Current density–voltage relationship for HEK293T cells (Ctrl, black) and ANO1-expressing cells (yellow), showing characteristic outward rectification for ANO1. **c**, Mean whole-cell current density traces elicited by 500 ms voltage ramps in cells expressing rEstus2s (Ctrl) or rEstus2s-T2A-K_Ca_3.1, using the same solutions as in b. **d**, Mean normalized *F–V* relationships of cells stably expressing rEstus2s (Ctrl) or the indicated co-expression constructs. **e**, Fluorescence traces of cells stably expressing rEstus2s (Ctrl) or one of the co-expression constructs. **f, g**, Minimum (f) and maximum (g) fluorescence excursions during recordings for 60 s (as in e). **h**, Mean fluorescence power spectra for cells expressing the indicated GEVIs. Sem is indicated by shading. **i, j**, Peak-to-peak fluorescence excursions (i) and absolute mean *V*_m_ (j) of individual and confluent cells expressing rEstus (Ctrl) or co-expression constructs. **k, l**, Minimum (k) and maximum (l) fluorescence excursions in cells expressing rEstus2s and either ANO1 or K_Ca_3.1 with or without 10 µM of the specific ion channel inhibitors Ani9 (ANO1) or Senicapoc (K_Ca_3.1). **m, n**, Mean change in *F*_green_ of rEstus (m) and rEstus2s (n) without (Ctrl) and with co-expression of the indicated channels, induced by the application of the Ca^2+^ ionophore ionomycin (200 nM). The number of analyzed cells is indicated in parentheses.

Individual HEK293T cells expressing only rEstus or rEstus2s exhibited spontaneous *V*_m_ fluctuations (Figs. 2e-g), as previously reported.^31^ The relative fluorescence change (peak-to-peak) was ≈55% higher for rEstus2s compared to rEstus, which is in agreement with the calculated ≈60% increase in sensitivity (d*F*/*F* d*V*^-1^) at -50 mV (the average resting *V*_m_ of HEK293T cells) from electrophysiological experiments (Supplementary Fig. S2). The spectral density of fluorescence recordings from rEstus2s was 2.4-fold higher than that of rEstus (Fig. 2h), underscoring its higher sensitivity. Expression of ANO1 or K_Ca_3.1 resulted in spontaneous depolarizing or hyperpolarizing events, respectively (Figs. 2e-g). The amplitudes of these events exceeded the endogenous excursions observed in HEK293T cells, and rEstus2s yielded higher relative fluorescence changes than rEstus. K_Ca_3.1 events had an average amplitude of 22.8 ± 1.1% (rEstus2s), a frequency of 2.0 ± 0.23 min^−1^, and a duration of 1.14 ± 0.12 s. ANO1 events had an average amplitude of -25.1 ± 0.9% and occurred with a frequency of 1.58 ± 0.21 min^−1^ (Fig. 2g). These depolarizing events of 0.58 ± 0.05 s were approximately half as long as the hyperpolarizing events induced by K_Ca_3.1. Importantly, these large *V*_m_ fluctuations were only detectable in isolated HEK293T cells and not in cells that were part of confluent cell layers (Fig. 2i).

Confluent and individual HEK293T cells exhibited similar resting *V*_m_ of about -50 mV (Fig. 2j). The expression of ANO1 depolarized the cells to about -30 mV, both in individual and confluent states. Remarkably, while individual cells overexpressing K_Ca_3.1 displayed similar or slightly lower resting *V*_m_ (≈ -40 mV) than control HEK293T cells, confluent K_Ca_3.1-expressing cells were hyperpolarized to about -65 mV. Since both channel types affected the resting *V*_m_, it is not necessarily clear that the spontaneous *V*_m_ excursions originate from the respective channel activity. However, acute application of the channel-specific inhibitors Ani9 (ANO1) and Senicapoc (K_Ca_3.1) eliminated these events, thus indicating channel-dependent activity (Figs. 2k,l). The responsiveness of the channels to intracellular Ca^2+^ level changes and the consequence for the average *V*_m_ was examined by application of the Ca^2+^ ionophore ionomycin, which is expected to maximally activate ANO1 and K_Ca_3.1 (Figs. 2m,n). Control HEK293T cells exhibited a transient hyperpolarization following ionomycin treatment. In K_Ca_3.1-expressing cells, this response was markedly enhanced and sustained, whereas ANO1-expressing cells displayed pronounced depolarization. Ionomycin-induced fluorescence changes were larger in cells expressing rEstus2s than rEstus, reflecting the increased voltage sensitivity of rEstus2s.

These results indicate that the *V*_m_ fluctuations arise from spontaneous openings of ANO1 and K_Ca_3.1 channels and suggest that, in cells expressing these oncochannels, the resting *V*_m_ is not clamped near the respective Cl^−^ or K^+^ Nernst potentials.

### Ca^2+^ remains cellularly restricted while *V*_m_ changes spread through open gap junctions

ANO1 and K_Ca_3.1 channels induce spontaneous changes in *V*_m_ in HEK293T cells (Fig. 2), and are activated by elevations in intracellular Ca^2+^. To investigate whether channel activity correlates with transient increases in intracellular Ca^2+^, we used the red fluorescent Ca^2+^ indicator K-GECO alongside rEstus2s and performed simultaneous live-cell imaging of *V*_m_ and Ca^2+^ (Fig. 3a).

**Figure 3.**
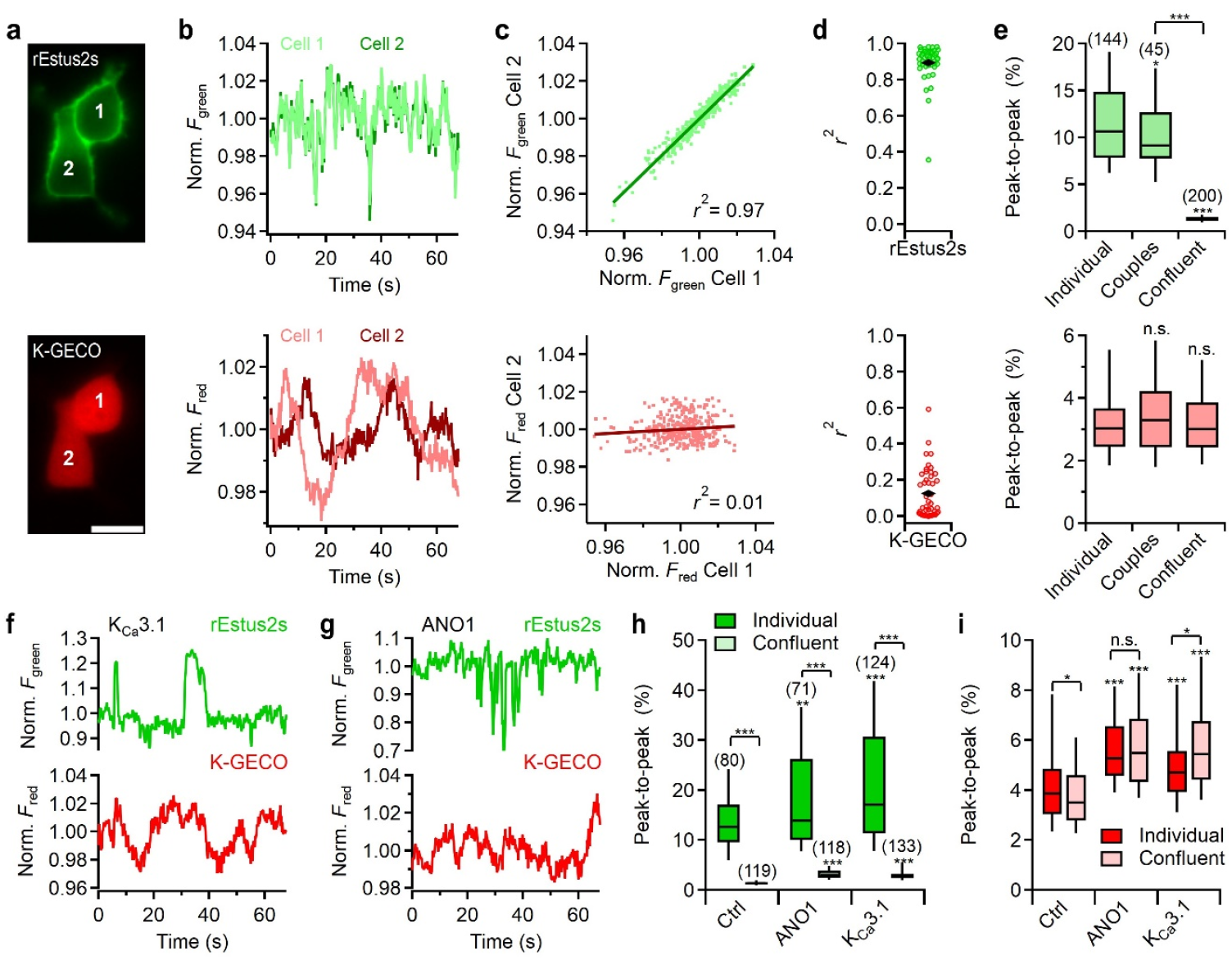
The fluctuations in *V*_m_, but not in Ca^2+^, correlate with the size of electrically connected cell networks. **a**, Fluorescence images of a pair of HEK293T cells stably expressing K-GECO-T2A-rEstus2s. Scale bar, 20 µm. top, *F*_green_; bottom, *F*_red_. **b**, Representative fluorescence traces of rEstus2s (top) and K-GECO (bottom) from the cells in a, indicating spontaneous changes *V*_m_ and intracellular free Ca^2+^, respectively. Traces were recorded with alternating excitation at 470 nm and 530 nm. **c**, Linear correlation of the time traces for rEstus2s (top) and K-GECO (bottom) from the cell pair shown in a. Coefficients of determination (r^2^) for the linear correlations (bold lines) are indicated. **d**, r^2^ values from linear correlations of rEstus2s and K-GECO fluorescence for 45 cell pairs. Black rhombs are means. **e**, Peak-to-peak excursions of relative rEstus2s (top) and K-GECO (bottom) fluorescence in individual cells, cell pairs (as in d), and confluent cells. **f, g**, Representative normalized fluorescence traces of rEstus2s (top) and K-GECO (bottom) in HEK293T cells stably expressing K-GECO-T2A-rEstus2s-T2A-K_Ca_3.1 (f) or K-GECO-T2A-rEstus2s-T2A-ANO1 (g). **h, i**, Peak-to-peak excursions of relative rEstus2s (h) and K-GECO (i) fluorescence in individual and confluent cells expressing K-GECO-T2A-rEstus2s (Ctrl) or additionally one of the indicated ion channel types. The number of analyzed cells is indicated in parentheses in h.

In HEK293T cells stably expressing K-GECO-T2A-rEstus2s, spontaneous fluctuations in both intracellular Ca^2+^ and *V*_m_ were observed (Fig. 3b). Measurement of spontaneous *V*_m_ fluctuations in neighboring cells can be used to estimate the electrical coupling between them because strongly correlated *V*_m_ indicates electrical connection with low resistance. We thus quantified electrical cell connectivity by fluorescence fluctuation correlation analysis (FFCA)^31^ applied to pairs of neighboring cells (Figs. 3b,c). Cell pairs exhibited strong synchrony in *V*_m_ but weak synchrony in Ca^2+^ signals (Fig. 3d), indicating that *V*_m_ changes propagate efficiently between neighboring HEK293T cells, whereas Ca^2+^ changes remain locally confined. Consistent with this observation, the amplitudes of *V*_m_ fluctuations but not Ca^2+^ fluctuations were strongly attenuated in confluent cell layers (Fig. 3e). If suppression of *V*_m_ fluctuations in confluent cultures were primarily driven by physical cell-cell contact and activation of membrane surface receptor pathways, a substantial reduction in *V*_m_ volatility would be expected upon formation of cell pairs. However, *V*_m_ fluctuations in cell pairs more closely resembled those observed in isolated cells rather than those in confluent cultures, indicating that the physical cell-cell contact alone has only a marginal effect on *V*_m_ volatility and is unlikely to account for the pronounced suppression of *V*_m_ dynamics in confluent monolayers.

*V*_m_ excursions arising from Ca^2+^-activated ion channel activity were not always accompanied by changes in global intracellular Ca^2+^ levels (Figs. 3f,g). In addition, *V*_m_ fluctuations were strongly attenuated at high cell densities (Fig. 3h), whereas global Ca^2+^ levels were not significantly affected by confluency (Fig. 3i). Yet, intracellular Ca^2+^ was slightly more variable in cells expressing ANO1 or K_Ca_3.1 compared to control HEK293T cells. To test whether the activity of ANO1 and K_Ca_3.1 is, in fact, stimulated by intracellular Ca^2+^, cells were pretreated with the membrane-permeable Ca^2+^ chelator BAPTA-AM. BAPTA-AM markedly reduced depolarizing events associated with ANO1 and hyperpolarizing events associated with K_Ca_3.1 (Supplementary Fig. S8). These findings indicate that spontaneous ANO1 and K_Ca_3.1 activity is driven by local rather than global increases in intracellular Ca^2+^.

### Connexins mediate cell density-dependent attenuation of spontaneous electrical activity through gap junction coupling

Endogenous *V*_m_ changes in HEK293T cells are markedly attenuated with increasing cell density. This effect was evident regardless of the overexpression of ANO1 or K_Ca_3.1. A plausible explanation is passive low-pass filtering arising from the electrical coupling, presumably through gap junctions formed by connexins.

HEK293T cells, which we have shown above (Fig. 3d) exhibit efficient electrical coupling, express two major connexins: Cx43 (*GJA1*) and Cx45 (*GJC1*).^20^ We thus generated a Cx43 knockout (Cx43 KO) and a Cx43/Cx45 double knockout (Cx43/45 KO) HEK293T cell line using CRISPR-Cas9 (Fig. 4a) and confirmed the absence of the respective proteins by Western blot (Supplementary Fig. S7). Upon stable expression of rEstus2s in both knockout cell lines, FFCA was applied to quantify electrical coupling between neighboring cells (Figs. 4b-e). Similar to control HEK293T, Cx43 KO cells exhibited strong electrical coupling, whereas the coupling of Cx43/45 KO cells was strongly reduced (Fig. 4d). Overexpression of Cx43 in Cx43/45 KO cells restored strong electrical coupling, indicating that both Cx43 and Cx45 are functionally active in HEK293T cells and together account for most of the electrical coupling.

**Figure 4.**
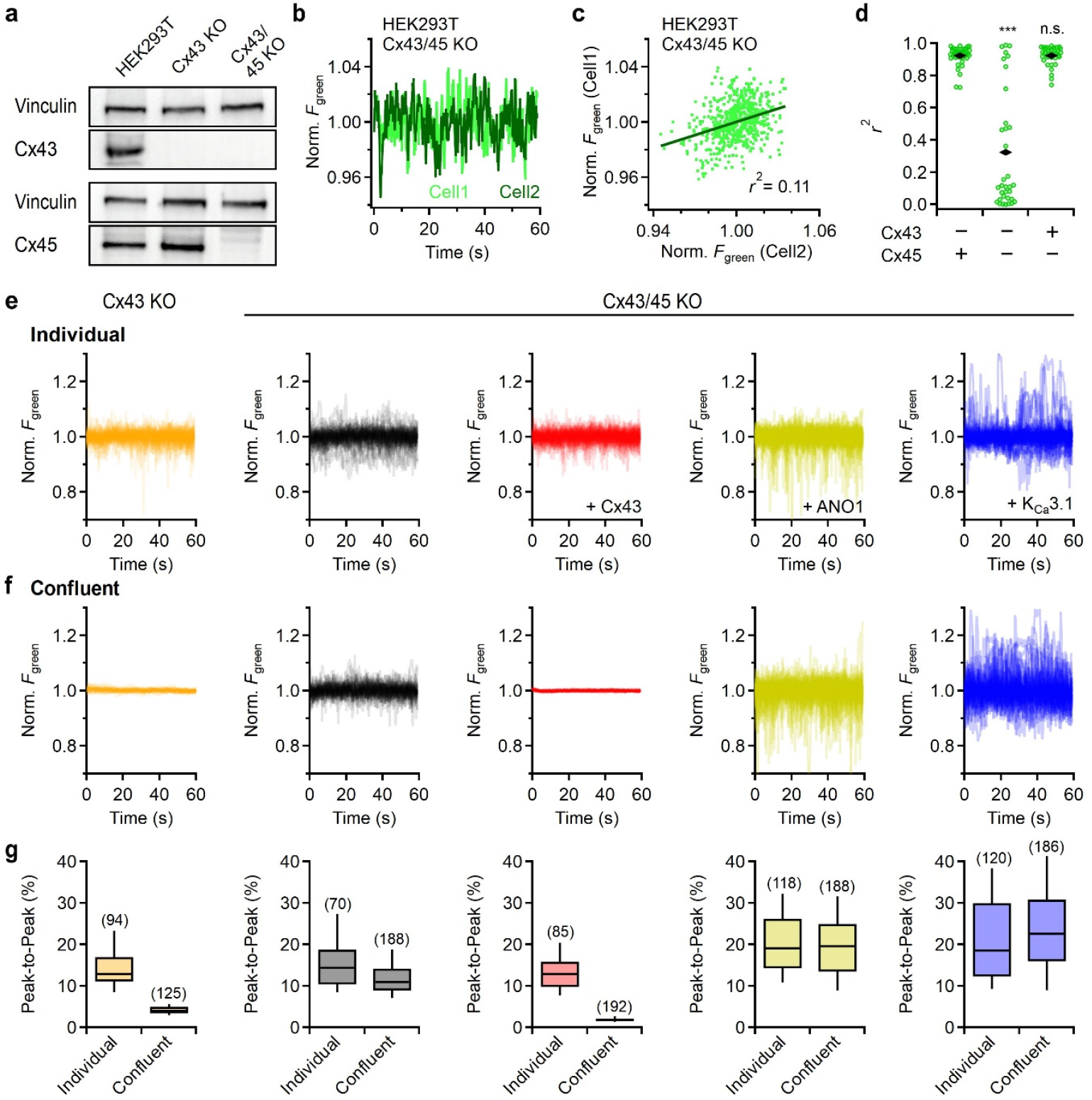
Connexin knockout abolishes cell density-dependent bioelectric filtering in HEK293T cells. **a**, Western blots of HEK293T cells and of Cx43 and Cx43/Cx45 knockout cell lines generated with CRISPR-Cas9. Vinculin was used as a loading control. **b**, Representative normalized fluorescence traces of two visually connected Cx43/Cx45 KO cells stably expressing rEstus2s. **c**, Linear correlation of the time traces from b with superimposed linear fit (bold line). **d**, r^2^ values from linear correlations of rEstus2s for Cx43 KO, Cx43/Cx45 KO, and Cx43/Cx45 KO with stable overexpression of Cx43. Black rhombs are means. **e, f**, Superimposed normalized fluorescence traces of 50 Cx43 KO cells each with rEstus2s, or Cx43/Cx45 KO cells with rEstus2s either alone or with additional overexpression of Cx43, ANO1, or K_Ca_3.1. Cells were either individual (e) or in a confluent layer (f). **g**, Peak-to-peak excursions of relative rEstus2s fluorescence in individual and confluent cells. The number of analyzed cells is indicated in parentheses.

*V*_m_ fluctuations in Cx43 KO cells were markedly diminished when cell cultures had reached confluency (Figs. 4e-g). In contrast, when both connexins were knocked out, the density-dependent reduction in *V*_m_ fluctuations was strongly reduced. Reintroduction of Cx43 into Cx43/45 KO cells fully restored the density-dependent attenuation of *V*_m_ fluctuations. Cx43/45 KO cells also expressing ANO1 or K_Ca_3.1 exhibited strong spontaneous *V*_m_ fluctuations as in regular HEK293T cells (Figs. 2i, 3h), but there was no longer any density-dependent attenuation of these fluctuations (Figs. 4e-g, Video 1). These data demonstrate that the observed attenuation of *V*_m_ excursions by ANO1 and K_Ca_3.1 (Fig. 3h) was also caused by gap junction coupling rather than surface receptor interaction.

### Loss of electrical coupling impairs bioelectric contact inhibition in cancer cell lines

Loss of gap junction coupling is an established phenomenon in early tumorigenesis, whereas the re-expression of connexins is often observed in later metastatic stages.^24^ Based on experiments with the Cx43/45 KO HEK293T cells (Fig. 4), we hypothesized that cell types lacking electrical coupling also lack density-dependent attenuation of electrical activity, a phenomenon we term bioelectric contact inhibition (BCI). We therefore examined a panel of cancerous and non-cancerous cell lines. We introduced rEstus2s into MCF-7 (breast cancer), Panc-1 (pancreatic ductal adenocarcinoma), A375 (melanoma), and non-cancerous HaCaT keratinocytes and HEK293T cells.

We characterized the expression of relevant ion channels and connexins using qPCR (Fig. 5a). mRNA coding for K_Ca_3.1 was highest in A375 cells and negligible in HEK293T cells. ANO1 was strongly expressed in HaCaT cells. Panc-1 cells predominantly expressed Cx45, whereas HaCaT cells primarily expressed Cx31 (*GJB3*). Transcript levels for all three tested connexins were low in MCF-7 cells, aligning with the weak electrical coupling among MCF-7 cells, as we have reported previously.^31^ The expression profiles are consistent with data from the Human Protein Atlas.^20^

**Figure 5.**
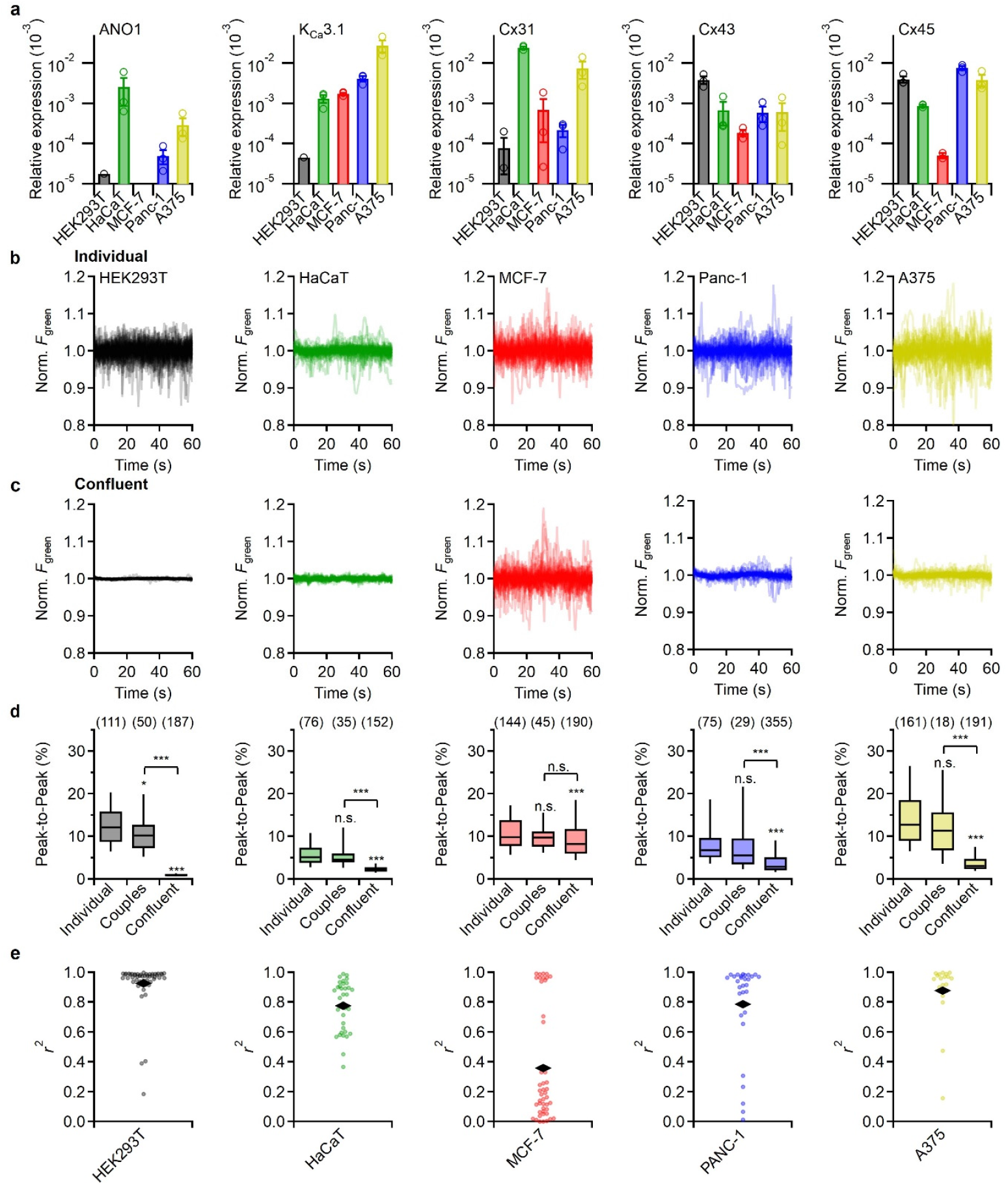
Cell types with reduced gap junction coupling lack cell density-dependent bioelectric filtering. **a**, Relative mRNA expression of the indicated ion channel and connexin genes in HEK293T, HaCaT, MCF-7, Panc-1, and A375 cells. Expression levels were determined by qRT-PCR and normalized to β actin. Data are means ± SEM. qPCRs were performed from three separate cDNA preparations. **b, c**, Superimposed normalized fluorescence traces (50 cells each) of individual (**b**) and confluent (**c**) cells stably expressing rEstus2s. **d**, Peak-to-peak excursions of relative rEstus2s fluorescence in individual cells, cell couples, and confluent monolayers. The number of analyzed cells is indicated in parentheses. **e**, Quantification of electrical coupling strength using Fluorescence Fluctuation Correlation Analysis on cell pairs. The coefficient of determination (r^2^) serves as a measure of synchrony. Black rhombs indicate the means. The number of analyzed cells is indicated in parentheses.

Individual cells of all cell lines displayed spontaneous *V*_m_ fluctuations (Figs. 5b-d). While HEK293T cells predominantly showed spontaneous depolarizations, Panc-1, MCF-7, and A375 cells exhibited a complex mix of depolarizing and hyperpolarizing events. This heterogeneity suggests that, beyond ANO1 and K_Ca_3.1, the electrical phenotype likely involves a diverse repertoire of other ion channels.

In confluent cell layers, all cell lines, with the exception of MCF-7, showed a significant reduction in *V*_m_ dynamics compared to individual cells or cell pairs (Figs. 5c,d), where the degree of reduction correlated with electrical coupling strength. FFCA (Fig. 5e) confirmed strong coupling in A375, HaCaT, and Panc-1 cells, while MCF-7 cells exhibited a substantially lower coupling strength. MCF-7 cells remained electrically active even at high density, with the majority of the monolayer functioning as uncoupled units. A375 cells, which had high *V*_m_ volatility in isolation, demonstrated a marked reduction in activity upon reaching confluency, indicative of strong BCI. It is worth noting that the electrical coupling in the other cancer lines was not always absolute. In Panc-1 monolayers, we observed subpopulations: some cells were fully decoupled, while others formed electrically coupled subsystems that generated slow oscillations independent of the surrounding monolayer (Video 2). This phenotype was notably more complex than that of HEK293T or HaCaT cells, where electrical activity was almost completely abolished at confluency. These data confirm that BCI is a general phenomenon of non-excitable cells that strictly depends on the electrical coupling strength within multicellular networks.

### Electrically coupled cells attenuate spontaneous *V*_m_ changes depending on the network size

The transition from individual cells to an electrically coupled syncytium can be described by an equivalent circuit model (Fig. 6a).^49^ In this model, the plasma membrane acts as a capacitor (*C*_m_) that resists *V*_m_ changes induced by current flow through stereotypic depolarizing (*R*_d_) and hyperpolarizing (*R*_h_) ion channels. *V*_m_ is determined by the open probability of these channels and the respective reversal potentials (*E*_rev_). When individual cells are connected through gap junction-mediated contacts, they establish low-resistance bridges (*R*_gj_) between the individual cellular circuits. For the specific case where *R*_d_≈*R*_h_≫*R*_gj_, the system behaves as a single large unit, where the total capacitance and conductance represent the sum of the individual cell components. Under this condition, the *V*_m_ of a single cell within the network becomes equivalent to the average *V*_m_ of the entire system. Conversely, the loss of connexins causes each cell to behave as an isolated electrical circuit, even at high cell density (Fig. 6b).

**Figure 6.**
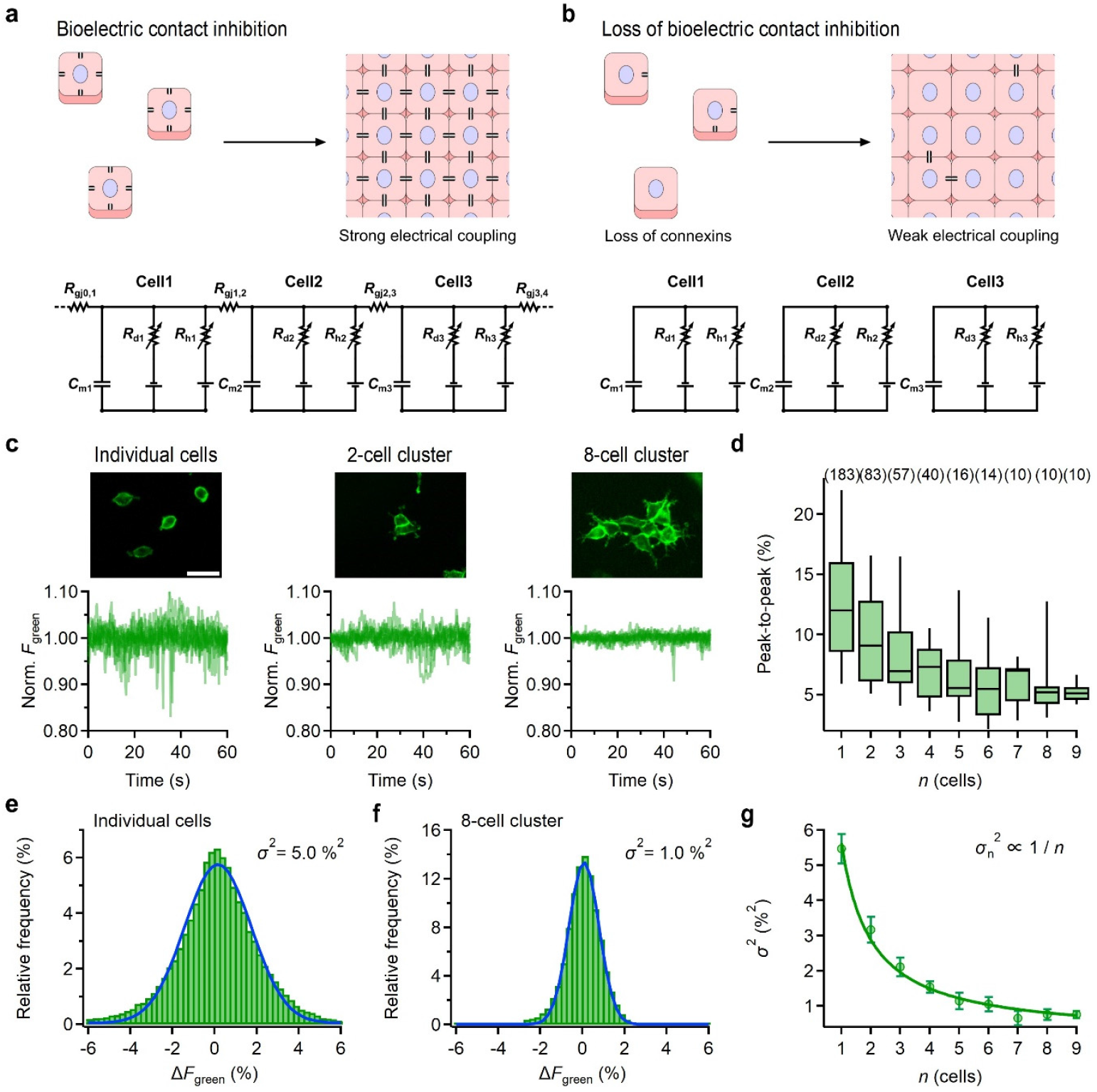
Electrically coupled cell networks attenuate *V*_m_ dynamics in individual cells depending on the network size. **a, b**, Schematic representation of bioelectric contact inhibition (a) and its loss (b). In multicellular networks, gap junctions (connexins) create strong electrical coupling, integrating individual cells into a syncytium. Loss of connexins, an early hallmark of tumorigenesis, results in weak or loss of electrical coupling. The electrical properties are depicted by the equivalent electrical circuit models: *R*_d_ and *R*_h_ are membrane resistances arising from stereotypical depolarizing and hyperpolarizing ion channels, respectively; *C*_m_ is the membrane capacitance; and *R*_gj_ the gap junction resistance. **c**, Top: Representative fluorescence images of HEK293T cells expressing rEstus2s in clusters of varying sizes (individual, 2-cell, and 8-cell clusters). Scale bar, 40 µm. Bottom: Superimposed normalized fluorescence traces of 10 individual cells or 10 clusters of the corresponding size. **d**, Peak-to-peak excursions of relative fluorescence as a function of cluster size (*n*). **e, f**, Average relative frequency distribution of fluorescence fluctuations for individual cells (e) and 8-cell clusters (f). Blue lines represent Gaussian fits. The variance (*σ*^2^) for each group is indicated, showing a reduction in volatility as cluster size increases. **g**, Variance of the normalized fluorescence as a function of cluster size. The solid line is the result of a 1/*n* fit (Eq. 4). The number of analyzed cells is indicated in parentheses.

Non-excitable cells are often characterized by low ion channel expression. When the open probability of these channels is low, they operate stochastically and act as independent noise sources that randomly inject current, causing spontaneous *V*_m_ fluctuations.^50^ In this state, *V*_m_ of an individual cell is highly sensitive to perturbations from even minor current injections. However, as cells electrically connect, both the total number of channels and the membrane capacitance scale linearly with the number of cells (*n*), thereby stabilizing the system. If the average resting *V*_m_ is maintained by only a few active channels, *V*_m_ volatility is expected to follow the law of large numbers. Assuming every cell is electrically similar, the variance in *V*_m_ (*σ*^2^) should be inversely proportional to the number of cells (*n*); hence, *σ*_n_^2^∝1/*n*.

To validate this model, we analyzed spontaneous *V*_m_ fluctuations in rEstus2s expressing HEK293T clusters of varying sizes (Fig. 6c). We found that the average peak-to-peak amplitude in *F*_green_ of these fluctuations systematically decreased with every additional cell added to the cluster (Fig. 6d). The *V*_m_ distributions around the mean were well-described by Gaussian functions (Fig. 6e), and the variance of an 8-cell cluster (Fig. 6f) was markedly reduced compared to that of individual cells. Notably, the reduction in *V*_m_ variance as a function of cluster size followed the predicted 1/*n* relationship (Fig. 6g). This confirms that *V*_m_ is maintained by stochastic channel fluctuations and shifts from stochastic to deterministic behavior depending on the network size. These findings establish that BCI is a fundamental physical property of electrically coupled networks, distinct from mechanical or chemical signaling mechanisms.

## Discussion

One central achievement of this study is the development of rEstus2s. Current high-performing GEVIs are predominantly developed by directed evolution, which often requires hundreds to thousands of mutants to be tested on large-scale screening platforms.^36–40^ Here, we pursued a rational engineering strategy that combines knowledge from decades of fluorescent protein optimization with our systematic analysis of how mutations in the VSD shape voltage-dependent optical responses of ASAP-type GEVIs.^31^ While brighter than ASAP3, rEstus2s is slightly dimmer than rEstus and JEDI-1P. However, this reduction in brightness is outweighed by the substantial gain in sensitivity and dynamic range. To our knowledge, rEstus2s is more sensitive than any currently available GEVI across the entire physiological resting *V*_m_ range (0 to -100 mV). rEstus2s avoids saturation even at the edges of the physiological resting *V*_m_ range, allowing for the confident resolution of *V*_m_ changes below 1 mV. This feature is vital because the exact resting voltage of non-excitable and tumor cells is often unknown *a priori*. While literature frequently proposes that tumor cells are in general depolarized (0 to -50 mV),^5,51^ using quantitative *V*_m_ imaging, we previously demonstrated that the average *V*_m_ of HEK293T, MCF-7, and A375 cells actually resides in the -40 mV to -60 mV range.^31,44,47^ A limitation of rEstus2s is that it exhibits reduced excitation at 400 nm relative to rEstus and therefore cannot be calibrated by dual-excitation ratiometry. This limitation is offset by its ability to support 400/470 nm co-illumination. This illumination scheme accelerates response kinetics without compromising sensitivity, likely by promoting voltage-dependent *cis-trans* chromophore isomerization through excitation of the protonated chromophore state.^48^

The development of rEstus2s was motivated by the need for tools capable of resolving how cancer-associated alterations, particularly the loss of connexin and the upregulation of oncochannels, reshape the electrical dynamics of non-excitable cells. While ion channels such as ANO1 and K_Ca_3.1 have been associated with enhanced ERK signaling and altered proliferative or migratory behavior, these relationships are typically inferred from long-term phenotypic assays.^13–20^ Here, we focus on the fast *V*_m_ dynamics generated by ion channel activity and demonstrate that these dynamics are fundamentally constrained by gap junction-mediated electrical coupling. Our results establish a unifying biophysical framework (BCI) describing how *V*_m_ fluctuations driven by ion channels are passively attenuated in electrically coupled multicellular networks, independent of potential downstream signaling pathways.

One of the prevailing models in cancer bioelectricity posits that a depolarized steady-state *V*_m_ is a key driver of tumorigenesis.^30,52,53^ We observed that while overexpression of the oncochannels K_Ca_3.1 and ANO1 shifted average *V*_m_ in opposite directions, *V*_m_ remained far from the reversal potentials for K^+^ or Cl^−^, respectively. This indicates that *V*_m_ is not dominated by any single ion species but instead reflects a balance among various conductances. The observation that the *V*_m_ variance scales with cluster size according to a 1/*n* relationship suggests that resting *V*_m_ is maintained by a small number of stochastically opening channels, rendering isolated cells or small clusters electrically volatile. Analogous to neuronal principles, where low channel density shifts a system from deterministic to stochastic behavior,^54,55^ non-excitable cells exhibit substantial *V*_m_ fluctuations at the single-cell level. Electrical coupling through gap junctions suppresses this volatility, acting as a biophysical noise filter.^35,56^ This noise filter is lost if gap junctions are downregulated during early tumorigenesis.^23,24^

Computational models by Cervera *et al*.^49^ demonstrate that multicellular networks can lock cells into synchronized, bistable resting *V*_m_ states. However, a critical distinction resides in the time domain: while these models primarily describe steady-state shifts occurring over hours, our data reveal that the electrical coupling through gap junctions suppresses *V*_m_ changes on the millisecond-to-second timescale.

Notably, despite robust electrical coupling through Cx43 and Cx45, intracellular Ca^2+^ dynamics remained unsynchronized even between neighboring HEK293T cells and were insensitive to network size. This suggests a functional separation of fast electrical coupling and slower metabolic or second-messenger coupling. Although connexins can support Ca^2+^ wave propagation indirectly via diffusion of signaling molecules such as IP_3_,^23^ strong intracellular buffering and limited permeability likely preclude rapid Ca^2+^ equilibration. In non-excitable contexts, gap junctions therefore appear to function primarily as low-resistance electrical conduits rather than as mediators of fast biochemical signaling.

Unlike contact inhibition of proliferation or locomotion, which describe biological outcomes, BCI refers to a biophysical constraint on *V*_m_ imposed by electrical coupling, which may act upstream of these processes.^10,57–59^ Mechanistically, while K_Ca_3.1 may enhance the driving force for Ca^2+^ entry and ANO1 may depolarize the membrane and e.g. directly activate ERK, the outcome is dictated by the syncytium. As long as a cell is embedded in a large coupled network, it is electrically clamped; it cannot significantly alter its *V*_m_ unless (1) its membrane conductance exceeds the junctional conductance, (2) it uncouples from the network, or (3) the network changes its *V*_m_ coherently. Thus, under healthy conditions, connexins act as a biophysical brake, forcing individual cells to conform to the resting *V*_m_ of the tissue.^49^

The scaling of *V*_m_ fluctuations with cluster size suggests a mechanism by which individual cells could infer network size.^1^ In this framework, localized current injection, for example through opening of K_Ca_3.1 or ANO1 channels in a single cell, could act as a rapid electrophysiological ping, with the resulting *V*_m_ amplitude providing immediate feedback about the size of the surrounding cellular network. In a growing tissue, this feedback could instruct proliferation until the network reaches a terminal size, at which point *V*_m_ signals are attenuated below a threshold required to sustain growth. It remains unresolved if endogenous *V*_m_ volatility itself serves as a signaling cue by transiently crossing critical voltage thresholds, or whether electrical stability primarily prevents individual cells from reaching such thresholds. These possibilities are not mutually exclusive and may differ among processes such as migration, proliferation, and differentiation. Because *V*_m_ is an emergent, system-level property rather than a purely cell-autonomous variable, it may also enable collective decision-making, permitting state transitions only when a sufficient fraction of the network changes coherently. Addressing these questions will require high-resolution, spatiotemporally resolved measurements linking *V*_m_ dynamics to downstream signaling events and lies beyond the scope of the present study. By providing both an ultrasensitive GEVI and a coherent biophysical framework, this study lays the groundwork for decoding the bioelectrical principles underlying tissue homeostasis and disease.

## Materials and Methods

### Molecular biology

For sensor development, an N-terminal fusion protein of the red fluorescent protein mKate2 and rEstus was used.^45^ Point mutations were introduced into mKate2-rEstus using megaprimer mutagenesis. The positions of mutations within the cpEGFP component of rEstus are indicated relative to the EGFP sequence. All constructs were cloned into a pCDNA3.1-puro vector, as previously described.^31^ All sequences were validated by Sanger sequencing (Eurofins).

For stable coexpression of ANO1, K_Ca_3.1, or Cx43, fusion proteins of rEstus/rEstus2s derivatives with the self-cleaving T2A peptide were generated. The promoter of pCDNA3.1-puro was replaced with an EF1α promoter. For stable expression with the Ca^2+^ indicator K-GECO,^61^ a vector containing one or two T2A peptides was generated in the following configurations: K-GECO-T2A-rEstus2s, K-GECO-T2A-rEstus2s-T2A-ANO1, or K-GECO-T2A-rEstus2s-T2A-K_Ca_3.1. For lentiviral production rEstus2s was cloned into pCDH-EF1α-Puro-CMV. All constructs were validated by sequencing. For detailed sequence information, see the attached files.

### Cell culture

HEK293T cells were obtained from the Centre for Applied Microbiology and Research (CAMR; Porton Down, Salisbury, UK). MCF-7 cells were obtained from the European Collection of Authenticated Cell Cultures (ECACC; Porton Down, Salisbury, UK). A375 and Panc-1 Cells were obtained from American Type Culture Collection (ATCC; Manassas, VA, USA). HaCaT cells were obtained from (Cytion; Heidelberg, Germany).

HEK293T and MCF-7 cells were cultured in DMEM/F-12 (Thermo Fisher Scientific), while A375, HaCaT, and Panc-1 cells were maintained in DMEM (Sigma-Aldrich). All media were supplemented with 10% fetal bovine serum (FBS). Cells were maintained at 37°C in a humidified incubator. A375 cells were kept at 10% CO_2_, while all other cell lines (HEK293T, MCF-7, HaCaT, and Panc-1) were maintained at 5% CO_2_.

For the generation of stable HEK293T cell lines, cells were transfected at sub-confluency in T25 culture flasks using Roti®-fect with 4 µg of rEstus or rEstus2s in pCDNA3.1-Puro, or with rEstus-T2A-ANO1, rEstus-T2A-K_Ca_3.1, rEstus2s-T2A-ANO1, rEstus2s-T2A-K_Ca_3.1, K-GECO-T2A-rEstus2s, K-GECO-T2A-rEstus2s-T2A-ANO1, or K-GECO-T2A-rEstus2s-T2A-K_Ca_3.1 in pCDNA3.1-EF1α-Puro. Cells were selected for two weeks in DMEM/F-12 supplemented with 10% FBS and 10 µg/ml puromycin (Gibco), with the medium refreshed every three days. Stable cell lines were subsequently maintained in medium without puromycin.

To generate stable cell lines, A375, and MCF-7 cells were transfected with the rEstus2s-pcDNA3.1(+)-Puro construct via electroporation and selected with 10 µM puromycin. Panc-1 and HaCaT cells were transduced using a lentiviral vector using a lentiviral vector (rEstus2s-pCDH-EF1 α -Puro) and selected with 4 µg/ml and 3 µg/ml puromycin, respectively. Polyclonal populations were enriched by fluorescence-activated cell sorting (FACS).

### Lentiviral production and transduction

Lentiviral particles were produced by transient co-transfection of HEK293T cells. One day prior to transfection, 3.2×10^6^ HEK293T cells were seeded into 60-mm culture dishes in high-glucose DMEM (Thermo Fisher Scientific) supplemented with 10% FBS and maintained at 37°C with 5% CO2. At 80-90% confluency, cells were transfected with 6 µg of pCDH-EF1α-Puro-CMV-rEstus2s and the packaging plasmids pMDLg/PRRE (3 µg; Addgene #12251), pRSVrev (2.5 µg; Addgene #12253), and pMD2.G (1.5 µg; Addgene #12259) using 25 µl Lipofectamine 2000 (Thermo Fisher Scientific) in Opti-MEM.

The culture medium was replaced 16 h after transfection. Virus-containing supernatants were harvested at 24 h and 48 h, filtered through a 0.45-µm cellulose acetate filter (Carl Roth), and stored at -80°C. For transduction, 1×10^5^ Panc-1 or HaCaT cells were seeded per well of a 6-well plate and incubated with 600 µl of lentiviral supernatant. After 24 h, the medium was replaced, and stable transformants were selected using puromycin (4 µg/ml for Panc-1; 3 µg/ml for HaCaT). Selection efficiency was monitored using non-transduced control cells.

### CRISPR-Cas9 knockout

For generating CRISPR-Cas9 knockout cell lines, we used the lentiCRISPRv2 system. The following guide RNAs (gRNAs) from the human CRISPR knockout pooled library A (GeCKOv2) were cloned into the lentiCRISPRv2 (Addgene #52961):^62^

*GJA1* (Cx43): 19135: TCAGCGCACCACTGGTCGCA, 19136: TGTGTTCTATGTGATGCGAA

*GJC1* (Cx45): 19177: CATCTTCCCGAATCCGTCGT, 19179: GCAAGCCCTATGCAATGCGC

HEK293T cells were transiently transfected with the lentiCRISPRv2 expression plasmids using Roti®-fect. One day after transfection, cells were selected for 3 days with 10 μg/ml puromycin to eliminate non-transfected cells. Following selection, individual clones were isolated by serial dilution into 96-well plates. Individual clonal cell lines were grown to confluency and maintained in a T-25 culture flask. Successful knockout was confirmed by Western blot analysis. The Cx43 single knockout cell line was generated first using gRNAs 19135 and 19136. Based on the Cx43 knockout (Cx43 KO) line, a Cx43/Cx45 double knockout (Cx43/45 KO) line was subsequently generated using gRNAs 19177 and 19179.

### Western Blot

Cells were washed once with ice-cold phosphate-buffered saline (PBS) and lysed in RIPA buffer (50 mM Tris pH 7, 150 mM NaCl, 1% Triton X-100, 0.1% SDS, 2 mM EGTA) supplemented with cOmplete protease inhibitor cocktail (Roche; distributed by Merck). Lysates were clarified by centrifugation (14,000 × g, 10 min, 4°C), and samples were incubated at 70°C for 10 min. Total protein (15 µg per lane) was resolved on 12% polyacrylamide gels alongside a size standard (PageRuler Prestained Protein Ladder, Thermo Fisher Scientific). Proteins were transferred to polyvinylidene difluoride (PVDF) membranes and blocked in Tris-buffered saline containing 0.1% Tween-20 (TBST) and 5% non-fat dry milk. Membranes were probed with mouse monoclonal antibodies from Santa Cruz Biotechnology: anti-Cx43 (sc-271837, 1:1000), anti-Cx45 (sc-374354, 1:1000), and anti-Vinculin (sc-73614, 1:2000). A peroxidase-conjugated goat anti-mouse secondary antibody (SeraCare, #5220-0341; 1:10000) was used for detection. Chemiluminescence was imaged using the Fusion FX Edge system with EvolutionCapt software (Vilber Lourmat).

### qPCR

Total RNA was isolated from confluent cells using the RNeasy Mini Kit (Qiagen; cat. no. 74104). cDNA was generated using the First Strand cDNA Synthesis Kit (Thermo Fisher Scientific; #K1612). qPCR was performed on a Mastercycler ep realplex (Eppendorf) using Maxima SYBR Green qPCR Master Mix (Thermo Fisher Scientific; #K0251). Primers for *GJC1* (Cx45; FH1/BH1-*GJC1*), *GJA1* (Cx43; FH1/BH1-*GJA1*), and *GJB3* (Cx31; FH1/BH1-*GJB3*) were obtained from Merck. Primers for *ANO1* (QT00076013), *KCNN4* (QT00003780), and β-actin (QT0095431) were obtained from Qiagen. All qPCR results were normalized to β-actin levels.

### Combined electrophysiology and fluorescence imaging

For electrophysiological recordings, HEK293T cells were plated on 35-mm glass-bottom dishes (Ibidi, Martinsried, Germany) and transfected the following day with 1 µg of plasmid DNA per dish using Roti®-fect (Carl Roth, Karlsruhe, Germany). Electrophysiological and imaging experiments were performed one or two days after transfection.

Electrophysiological recordings were performed using an EPC10 double patch-clamp amplifier controlled by PatchMaster software (HEKA Elektronik, Lambrecht, Germany) on an Axio Observer inverted microscope (Carl Zeiss, Jena, Germany). For fluorescence excitation, a Colibri-2 LED illumination system and an EC-Plan Neofluar 40× oil-immersion objective (NA 1.3, Zeiss) were used. Images were acquired with an ORCA-Flash 4.0 Digital Camera (C11440, Hamamatsu, Japan) controlled by SmartLux software (HEKA Elektronik).

For sensor development, mKate2 fusion constructs were used. mKate2 was excited with a 530-nm LED (Zeiss) that had an output of 1.8 mW. The filter set was as follows: excitation: BP 545/40 (Chroma), beam splitter: 565 LPXR (Chroma), emission: BP 608/65 (Delta Optical Thin Film). For excitation of GEVIs, a 470-nm LED (Zeiss) with an output of 0.7 mW was used. The filter set was as follows: excitation: BP 470/40 (Chroma), beam splitter: 495 LPXR (Chroma), emission: BP 525/50 (Delta Optical Thin Film). Images were acquired with a 90-ms camera exposure time at 5 Hz.

For limit of detection recordings (Figs. 2e-h), images were acquired with a 40 ms camera exposure time at 20 Hz. Excitation was achieved through simultaneous illumination with a 470-nm LED (Zeiss) with a BP 470/40 filter at 0.74 mW and a 400-nm LED (Zeiss) with a BP 400/10 filter (Thorlabs) at 0.12 mW. The filter cube contained a 495 LPXR beamsplitter and a BP 525/50 emission filter. The 400-nm LED was used to accelerate the sensor kinetics, as shown in Supplementary Fig. S4.

Sensor kinetics was evaluated using whole-cell patch clamp combined with photometry on an Axio Observer inverted microscope (Zeiss) equipped with an EC-Plan Neofluar 40× oil-immersion objective. Excitation light from a 470-nm LED (M470L1, Thorlabs) was passed through a modified filter set 38 (Zeiss), in which the excitation filter was replaced with a 492/SP filter (Semrock, Rochester, NY, USA). Emission light was detected using a photodiode (FDU photodiode with viewfinder, T.I.L.L. Photonics, Gräfelfing, Germany) and recorded at 20 kHz using PatchMaster software (HEKA Elektronik).

Patch-clamp pipettes were made from borosilicate glass, coated with dental wax, and fire-polished to obtain resistances of 1-2 MΩ. For all electrophysiological recordings, the bath solution contained 146 mM NaCl, 4 mM KCl, 2 mM CaCl_2_, 2 mM MgCl_2_, and 10 mM HEPES, adjusted to pH 7.4 at 23°C using NaOH. The pipette solution for sensor calibration contained 130 mM KCl, 2.5 mM MgCl_2_, 10 mM EGTA, and 10 mM HEPES, adjusted to pH 7.4 at 23°C using KOH. For recordings of K_Ca_3.1 and ANO1 currents, the pipette solution contained 1 mM NaCl, 70 mM K^+^-gluconate, 50 mM KCl, 8 mM CaCl_2_, 10 mM EGTA, 1.5 mM MgCl_2_, and 20 mM HEPES, adjusted to pH 7.4 at 23°C using KOH. The intracellular free Ca^2+^ concentration in this solution was calculated to be approximately 300 nM using WEBMAXC STANDARD.

### Live-cell fluorescence imaging

For experiments with low cell densities, cells were seeded at a density of 20,000 cells per 35 mm glass-bottom dish (Ibidi) and imaged 48 h after plating. For confluent culture experiments, cells were seeded at either 80,000 cells per dish (imaged after 72 h) or 160,000 cells per dish (imaged after 48 h).

For live-cell imaging experiments (Figs. 2-6), stably transfected cells were washed once and subsequently maintained in the external bath solutions used in electrophysiological experiments, supplemented with 5 mM glucose. Cells were equilibrated in this solution for 15 min prior to image acquisition. Recordings were limited to a maximum duration of 15 min per sample. All recordings were performed at ambient temperature (23°C).

Images were acquired at a frame rate of 5 Hz (Figs. 2,3) with alternating 90 ms exposures for excitation at either 400 nm (0.40 mW) and 470 nm (0.74 mW), or 470 nm (0.74 mW) and 530 nm (1.8 mW). Alternatively, images were acquired at 10 Hz (Figs. 3-5) using simultaneous illumination at 400 nm (0.12 mW) and 470 nm (0.74 mW). All experiments were performed using the same epifluorescence setup described for sensor calibration.

Cluster size analysis (Fig. 6) was performed on an inverted Eclipse Ti fluorescence microscope (Nikon, Tokyo, Japan) equipped with a 10× Plan Apo λ objective (NA 0.45, Nikon). Excitation was provided by an X-Cite 120 LED light source (Excelitas Technologies, Waltham, MA, USA) through a GFP-3035D-000 filter set (Semrock, Rochester, NY, USA). Images were acquired at a frame rate of 4 Hz using a 14-bit DS-Qi2 CMOS camera (Nikon) controlled by NIS-Elements 4.6 software (Nikon).

### Image data analysis

All raw image sets were analyzed using Fiji (ImageJ). For GEVI analysis, regions of interest (ROIs) were defined either along the cell periphery (Figs. 1-3) or over the entire cell area (Figs. 3-6). For FFCA and K-GECO experiments, ROIs were restricted to the cytosol to prevent signal contamination from the overlapping membranes of neighboring cells. All extracted data were background-corrected by subtracting the mean fluorescence of a cell-free region. Subsequent processing was performed in Igor Pro 9 (WaveMetrics). Initial transients, primarily attributed to photoswitching, were removed, and traces were corrected for photobleaching using a single-exponential decay function. FFCA was performed as described previously.^31^ For FFCA experiments and the experiments in Figs. 4-5, fluorescence traces were smoothed using a binning factor of 2.

To determine the relative molecular brightness during sensor development (Fig. 1), the green fluorescence intensity (*F*_green_, GEVI) was normalized to the red fluorescence intensity (*F*_red_, mKate2). The mean steady-state fluorescence ratio (*R*=*F*_green_/*F*_red_) was plotted as a function of membrane voltage (*V*_m_) and fitted with a Boltzmann function (Equation 1)

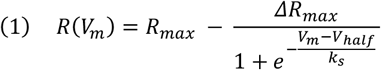

where *R*_max_ is the maximum fluorescence ratio, Δ*R*_max_ is the maximal change in fluorescence ratio, *V*_half_ is the voltage at half-maximal response, and *k*_s_ represents the slope factor. Fit parameters for all constructs are provided in Table S1.

The voltage sensitivity (brightness change per voltage unit) was described by the first derivative of Equation 1:

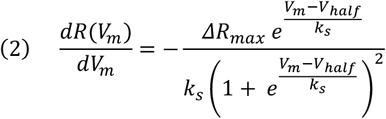

To calculate the fractional voltage sensitivity as a function of voltage, Equation 2 was divided by Equation 1.

For kinetic characterization of the GEVIs, the normalized fluorescence (*F*) response to voltage steps was analyzed using a double-exponential fit (Equation 3):

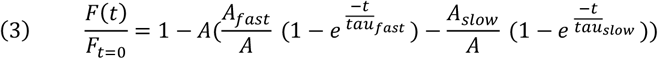

where *A* represents the amplitude and *τ* the time constant of the fast and slow components, respectively.

To characterize fluorescence fluctuations, we calculated the relative fluctuation magnitude normalized to the mean fluorescence intensity of each trace. Key parameters extracted included the peak-to-peak amplitude, maximum positive and negative excursions from the mean, and the time variance. For co-expression experiments with rEstus2s and ion channels, channel opening events were defined as fluctuations exceeding a specific threshold relative to the mean: a >10% increase for K_Ca_3.1 or a >15% decrease for ANO1.

The dependence of fluorescence volatility on cluster size (*n*) was modeled by fitting a reciprocal decay function to the *F*_green_ time variance (*σ*_n_^2^):

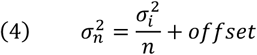

where *σ*_i_^2^ represents the variance of an isolated cell and the offset accounts for the intrinsic background noise floor of the measurement system. This model assumes that for small *V*_m_ fluctuations *F*_green_ is linearly proportional to *V*_m_.

### Statistics and reproducibility

Data are presented as means ± standard error of the mean (sem), as indicated in the figure legends. For box plots, the central line indicates the median, the box limits indicate the 25^th^ and 75^th^ percentiles (interquartile range), and the whiskers extend to the 10^th^ and 90^th^ percentiles. For datasets with small sample sizes, individual data points are overlaid. The number of analyzed cells or traces (n) is provided in parentheses within the Figures. For comparisons between two independent groups, a two-sided Wilcoxon rank-sum test was performed. For multiple comparisons against a control group (Fig. 1h), a one-way ANOVA followed by Dunnett’s post-hoc test was used. Statistical significance is indicated as follows: * p<0.05, ** p<0.01, *** p<0.001; n.s., not significant (p≥0.05). For imaging experiments, data were compiled from 3 to 6 independent culture dishes.

## Acknowledgments

We thank Angela Roßner, Silke Tonndorf-Martini, and Sassrika N. C. W. Dehiwalage for technical support. We are grateful to Julia Drube for assistance with CRISPR-Cas9 experiments and Ingrid Hilger for providing Panc-1 cells. We also thank Katrin Schubert and Michael Müller at the FACS core facility of the Fritz Lipmann Institute (FLI) for the sorting of stable cell lines.

## Author contributions

Conceptualization: PR

Methodology: PR, RH

Investigation: PR, RH, AGN, RS, RM, SR, KF

Formal Analysis: PR

Visualization: PR

Supervision: PR

Writing—original draft: PR

Writing—review & editing: PR, RH, AGN, RS, SHH, RM, SR

Funding Acquisition: SHH

## Data and materials availability

All data are available in the main text or the supplementary materials. The authors agree to make the data and materials supporting the results or analyses presented in the paper available upon reasonable request. The rEstus2s expression vector is available from Addgene (250991).

