## Supplementary Information for "Sub-Millivolt Voltage Imaging Reveals Gap Junction-Mediated Bioelectric Contact Inhibition"

**Supplementary Table 1**. Parameters of the fits according to Eq. 1 describing the brightness–voltage relationships of all constructs from Fig. 1.

| GEVI | n | $R_{max}$ | ±SEM | $\Delta R_{max}$ | ±SEM | $V_{half}$ | ±SEM | $k_s$ | ±SEM |
|---|---|---|---|---|---|---|---|---|---|
| **rEstus** | 5 | 2.07 | 0.05 | -1.60 | 0.03 | -41.1 | 2.8 | 28.5 | 0.3 |
| **ASAP3** | 5 | 1.15 | 0.04 | -0.85 | 0.02 | -60.0 | 2.9 | 40.2 | 0.9 |
| **JEDI-1P** | 6 | 1.68 | 0.07 | -1.23 | 0.06 | -35.6 | 2.3 | 21.2 | 0.5 |
| **rEstus-$(T65G)_{EGFP}$** | 5 | 0.65 | 0.04 | -0.58 | 0.03 | -27.9 | 1.3 | 25.2 | 0.6 |
| **rEstus-$(F46L)_{EGFP}$** | 5 | 1.99 | 0.03 | -1.56 | 0.03 | -22.4 | 1.2 | 27.0 | 0.7 |
| **rEstus-$(V68L)_{EGFP}$** | 5 | 0.25 | 0.02 | -0.18 | 0.02 | -29.7 | 2.7 | 29.0 | 0.8 |
| **rEstus-$(F46L\text{-}T65G)_{EGFP}$** | 5 | 0.90 | 0.03 | -0.81 | 0.03 | -24.5 | 0.7 | 24.71 | 0.18 |
| **rEstus-$(T65G\text{-}V68L)_{EGFP}$** | 5 | 1.02 | 0.06 | -0.93 | 0.06 | -24.2 | 1.5 | 24.6 | 0.3 |
| **rEstus-$(F46L\text{-}T65G\text{-}V68L)_{EGFP}$** | 5 | 1.31 | 0.03 | -1.20 | 0.04 | -23.0 | 2.5 | 25.2 | 0.4 |
| **rEstus2s (rEstus-G138N-T141I-$(F46L\text{-}T65G\text{-}V68L)_{EGFP}$)** | 5 | 1.24 | 0.06 | -1.17 | 0.06 | -58.3 | 1.3 | 30.0 | 0.24 |

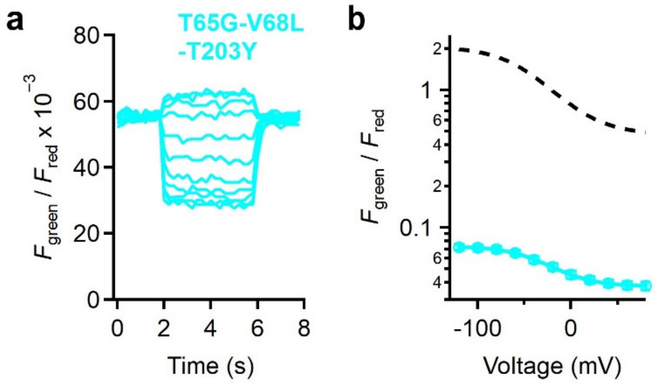


**Supplementary Figure S1. Fluorescence-voltage relationship of rEstus-T65G-V68L-T203Y.** **a**, Fluorescence response of rEstus-T65G-V68L-T203Y to membrane voltage changes. The green fluorescence of the GEVI was normalized to the red fluorescence of mKate2. Cells were held in the whole-cell voltage-clamp configuration. For the pulse protocol, see Fig. 1. **b**, Brightness-voltage relationship, derived from the steady-state fluorescence signal at the respective voltages. Data are means ± SEM, with superimposed Boltzmann-type fits (Eq. 1). The fit of rEstus is shown as a dark dashed line for reference.

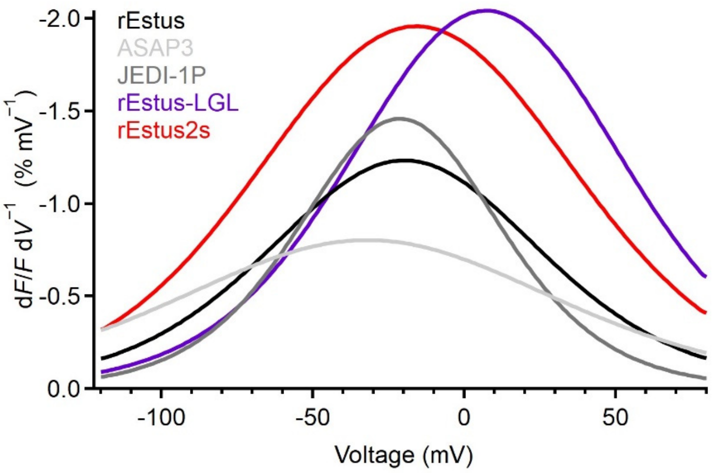


**Supplementary Figure S2. Voltage sensitivities of rEstus variants and established GEVIs.** The voltage sensitivity, defined as the fractional fluorescence change per unit voltage, is plotted as a function of $V_m$. Curves are the first derivative of the Boltzmann functions divided by the Boltzmann function fitted to the steady-state fluorescence–voltage relationships (Fig. 1e, f). rEstus2s (red) exhibits the highest sensitivity across the physiological resting $V_m$ range (-100 to 0 mV).

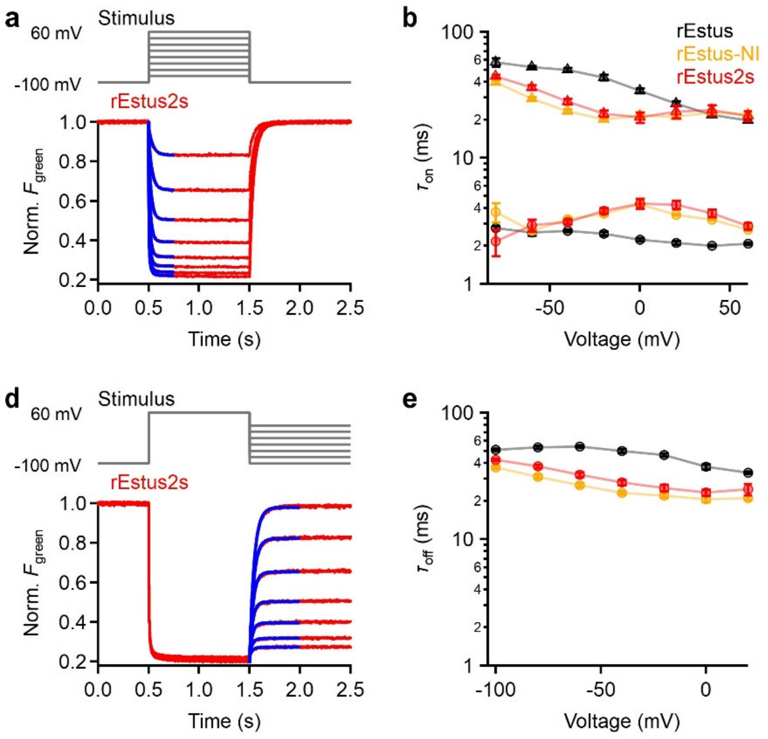


**Supplementary Figure S3. Voltage response kinetics of rEstus, rEstus2s, and rEstus-G138N-T141I. a,** Representative photometry measurement of the normalized $F_{green}$ from a HEK293T cell expressing rEstus2s, voltage-clamped according to the indicated protocol (top) at 23°C. The $F_{green}$ response to depolarization was fitted with a double-exponential function (blue, Eq. 2). **b,c,** Fast (circles) and slow (triangles) time constants (b) and relative amplitudes of the fast component (c), for rEstus, rEstus-G138N-T141I (rEstus-NI) and rEstus2s. **d,** $F_{green}$ response to repolarizing pulses with a superimposed single-exponential fit (blue). **e,** Time constants (τ) derived from the fit in d. Data are means ± SEM with n=5 for each construct. Straight lines connect data points in b, c, and e for clarity.

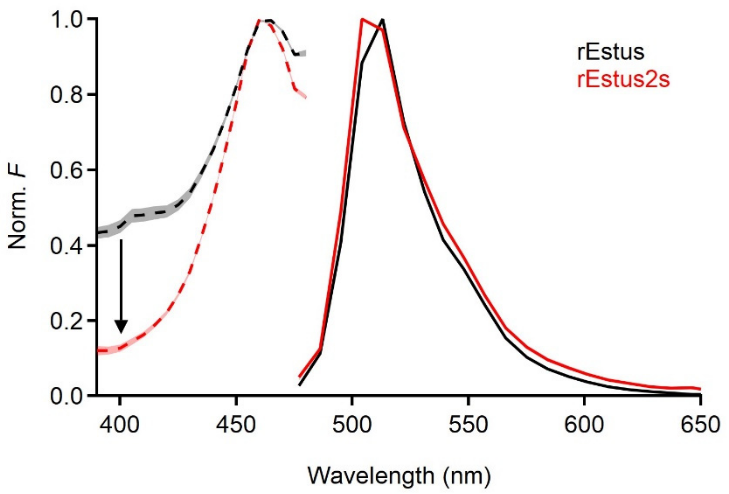


**Supplementary Figure S4. Excitation and emission spectra of rEstus and rEstus2s.** Mean normalized excitation (dashed lines) and emission (solid lines) spectra of rEstus (black) and rEstus2s (red). For the recording of emission spectra, the GEVIs were excited at 455 nm. Emission spectra were recorded on an LSM 980 (Carl Zeiss). Spectra were recorded from HEK293T cells without electrophysiological control. Note the loss of the excitation shoulder at around 400 nm for rEstus2s. Shaded areas represent SEM (n = 5-10 cells each).

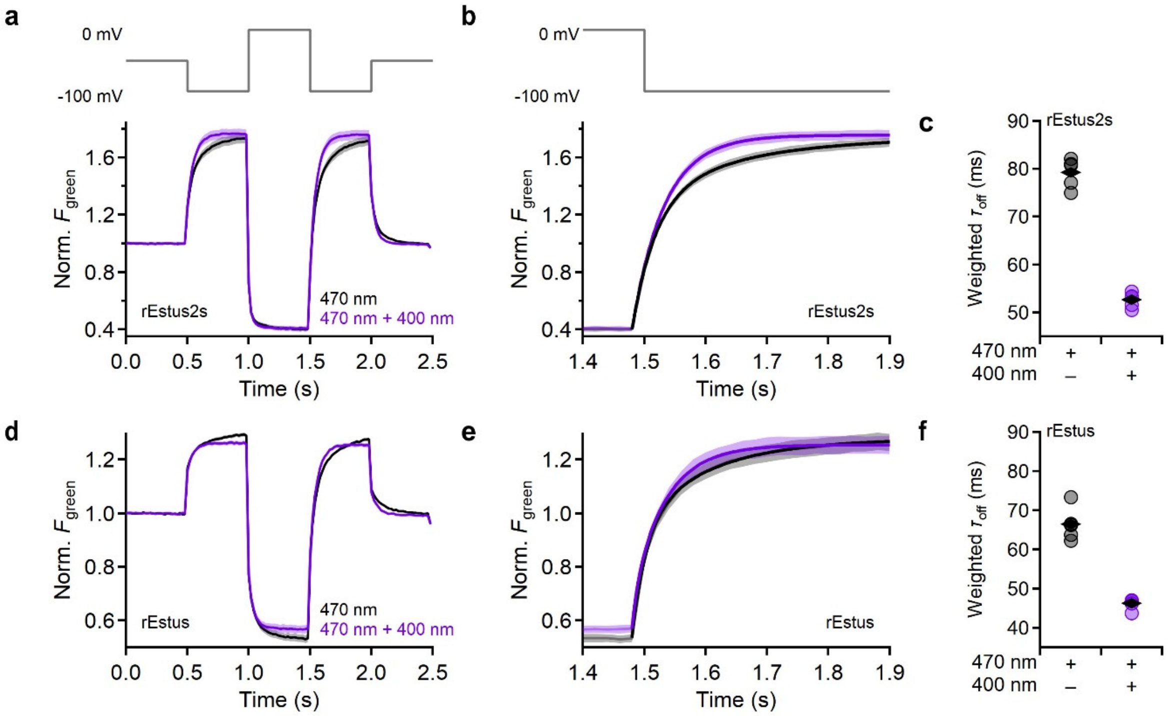


**Supplementary Figure S5. Excitation wavelength modulates the kinetics of rEstus and rEstus2s. a,** Normalized fluorescence response of a HEK293T cell expressing rEstus2s to the indicated pulse protocol (top) in whole-cell voltage clamp configuration at 23°C. Cells were illuminated with a 470 nm LED (black trace) or simultaneously with a 470 nm LED and a 400 nm LED (purple trace). Data are means; shading indicates SEM. **b,** Magnified section from panel a showing a repolarizing pulse from 0 to -100 mV with or without additional illumination at 400 nm. Traces were fitted with a double-exponential function. **c,** Amplitude-weighted time constants for a repolarizing pulse from 0 to -100 mV. Data points represent individual recordings; rhombs are means. **d-f,** As in a-c but for rEstus.

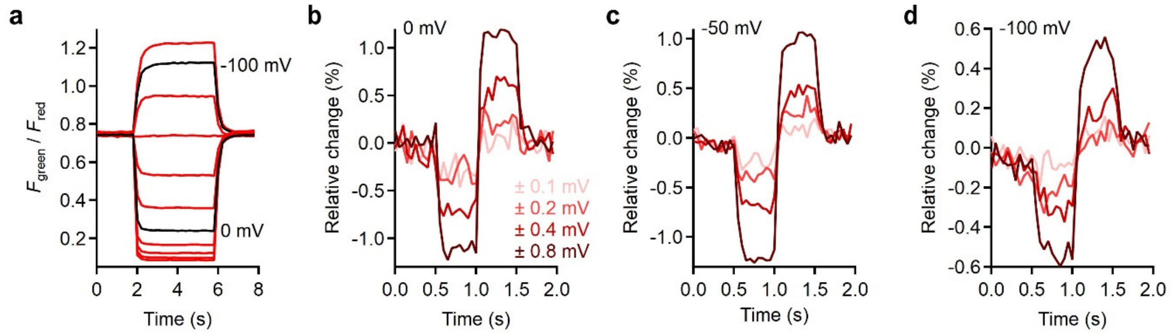


**Supplementary Figure S6. Detection of sub-millivolt changes with rEstus2s across the physiological resting $V_m$ range. a,** Superposition of representative normalized fluorescence traces ($F_{green}$ / $F_{red}$) of a HEK293T cell expressing mK2-rEstus2s, subjected to voltage steps between -120 and 80 mV from a holding potential of -60 mV. Notably, the fluorescence response does not saturate at the boundaries of the physiological resting $V_m$ range (-100 to 0 mV). The steady-state fluorescence–voltage relationship is shown in Fig. 1f. **b-d,** Representative fluorescence responses of rEstus2s to small bipolar square-wave voltage pulses with sub-mV amplitude. Traces were recorded at holding potentials of 0 mV (b), -50 mV (c), and -100 mV (d). Recordings in b-d were performed at 20 Hz with simultaneous 400 nm and 470 nm co-illumination to accelerate sensor kinetics.

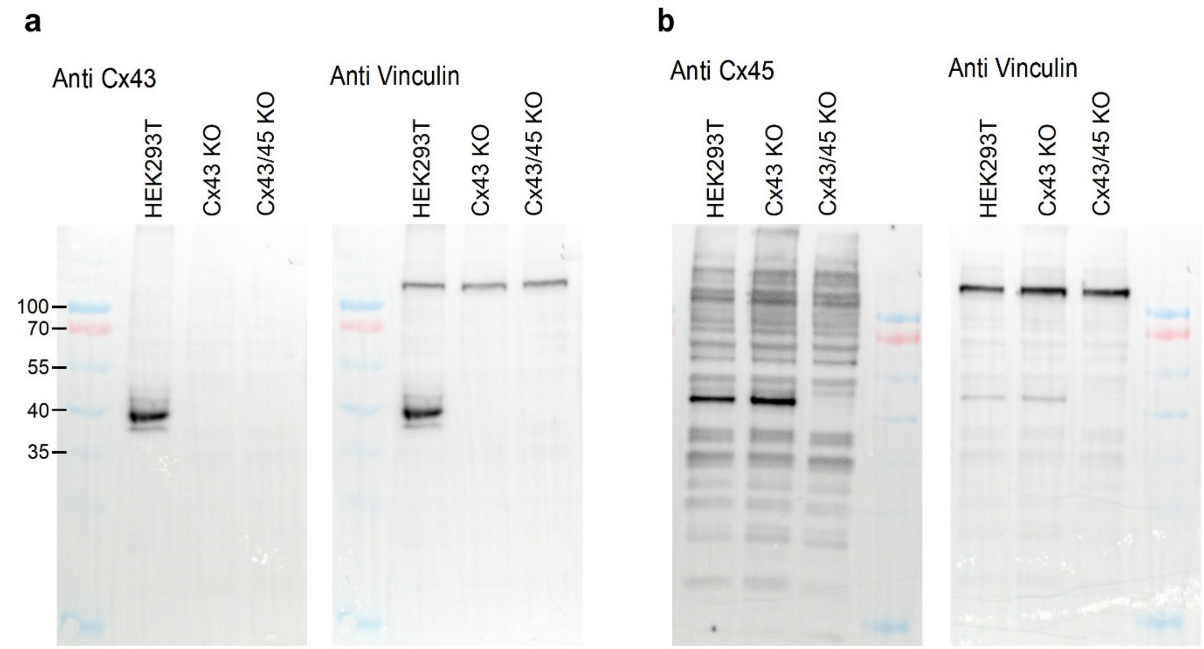


**Supplementary Figure S7. Validation of CRISPR-Cas9-mediated connexin knockout in HEK293T cells. a,** Western blot analysis of Cx43 expression in wild-type HEK293T cells, Cx43 knockout (Cx43 KO) cells, and Cx43/Cx45 double knockout (Cx43/45 KO) cells. Vinculin was used as a loading control. **b,** Western blot analysis of Cx45 expression in the indicated cell lines. Vinculin was used as a loading control.

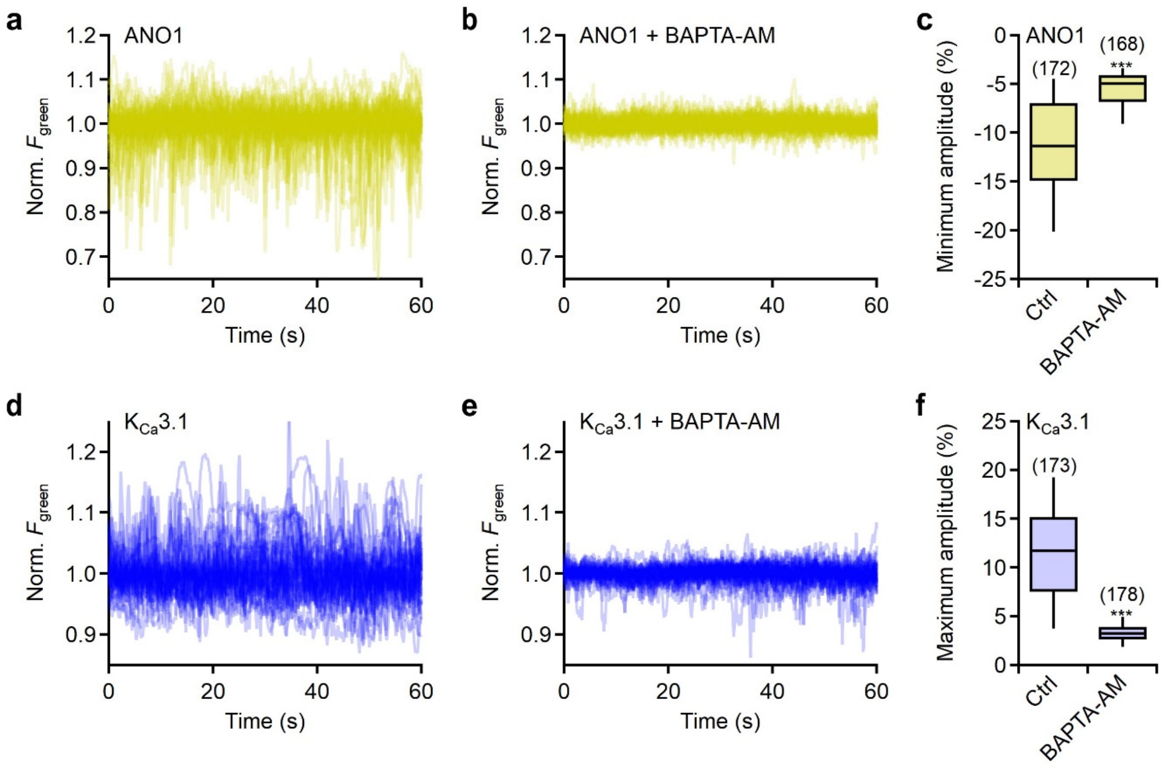


**Supplementary Figure S8. *$V_m$* fluctuations driven by $Ca^{2+}$-activated channels are suppressed by intracellular $Ca^{2+}$ buffering. a,** Superimposed normalized fluorescence traces of 50 confluent HEK293T Cx43/Cx45 KO cells stably expressing rEstus2s-T2A-ANO1. **b,** As in a but for cells pretreated for 1 h with 10 µM of the membrane permeable $Ca^{2+}$ chelator BAPTA-AM. **c,** Minimum fluorescence amplitudes of cells expressing rEstus2s and ANO1 with or without BAPTA-AM. The number of analyzed cells is given in parentheses. **d-e,** As in a-b but for HEK293T Cx43/Cx45 KO cells stably expressing rEstus2s-T2A-$K_{Ca}3.1$. **f,** Maximum fluorescence amplitudes of cells expressing rEstus2s and $K_{Ca}3.1$ with or without BAPTA-AM.
